# The wild species genome ancestry of domestic chickens

**DOI:** 10.1101/711366

**Authors:** Raman Akinyanju Lawal, Simon H. Martin, Koen Vanmechelen, Addie Vereijken, Pradeepa Silva, Raed Mahmoud Al-Atiyat, Riyadh Salah Aljumaah, Joram M. Mwacharo, Dong-Dong Wu, Ya-Ping Zhang, Paul M. Hocking, Jacqueline Smith, David Wragg, Olivier Hanotte

## Abstract

Hybridization and/or introgression play a key role in the evolutionary history of animal species. It is commonly observed in several orders in wild birds. The domestic chicken *Gallus gallus domesticus* is the commonest livestock species exploited for the benefit of mankind. The origin of its diversity remains unsettled. Here, we report a genome-wide analyses for signatures of introgression within domestic village chicken. We first established the genome-wide phylogeny and divergence time across the genus *Gallus*, showing the sister relationships between Grey junglefowl *G. sonneratii* and Ceylon junglefowl *G. lafayettii* and that the Green junglefowl is the first diverging lineage within the genus *Gallus*. Then, by analysing the whole-genome sequences of geographically diverse chicken populations, we reveal extensive bidirectional introgression between Grey junglefowl and domestic chicken and to a much less extent with Ceylon junglefowl. A single case of Green junglefowl *G. varius* introgression was identified. These introgressed regions include biological functions related to the control of gene expression. Our results show that while the Red junglefowl is the main ancestral species, introgressive hybridization episodes have impacted the genome and contributed to the diversity of domestic chicken, although likely at different level across its geographic range.

## Introduction

Despite the importance of domestic chicken *Gallus gallus domesticus* to human societies with more than 65 billion birds raised annually to produce meat by the commercial sector [1] and more than 80 million metric tons of egg produced annually for global human consumption, the origin and history of the genetic diversity of this major domesticate is only partly known. The Red junglefowl is the recognized maternal ancestor of domestic chicken [2, 3], with evidence from mitochondrial DNA (mtDNA) supporting multiple domestication centres [4] and the likely maternal contribution of several of its subspecies with the exception of *G. g. bankiva* (a subspecies with a geographic distribution restricted to Java, Bali and Sumatra).

However, the genus *Gallus* comprises three others wild species which may have contributed to the genetic background of domestic chicken. In South Asia, the Grey junglefowl *G. sonneratii* is found in Southwest India and the Ceylon junglefowl *G. lafayettii* in Sri Lanka. In South-East Asia, the Green junglefowl *G. varius* is endemic to Java and neighbouring islands [5] (Fig. 1*A*). Hybridization between the Red and the Grey junglefowls in their sympatric zones on the Indian subcontinent has been documented [5]. In captivity, hybridization between different *Gallus* species has been reported [6, 7], with Morejohn (1968) successfully producing F1 Red junglefowl x Grey junglefowl fertile hybrids in subsequent backcrossing with both species. Red junglefowl/domestic chicken mtDNA has been found in captive Grey junglefowls [8, 9] and the yellow skin phenotype is likely the result of the introgression of a Grey junglefowl chromosomal fragment into domestic chicken [10]. Captive F1 hybrids between female domestic chicken and male Green junglefowl, prized for their plumage colour and distinct voice, are common in Indonesia where they are known as Bekisar [5]. More generally, inter-species hybridization and introgression is an evolutionary process that plays a major role in the genetic history of species and their adaptation [11]. It may occur in the wild, when species live in sympatry, or in captivity following human intervention. While unravelling how it happens and detecting its signatures at the genome level is central to our understanding the speciation process, inter-species hybridizations are commonly practiced in agricultural plants and livestock for improving productivity [12] with hybridization also known to occur between domestic and wild species in several taxa [13]. Hybridization and introgression are relatively common in wild birds, including in Galliformes [6, 14-17]. For example, the genetic integrity of the rock partridge *Alectoris graeca* is being threatened in its natural habitat through hybridization with the introduced red-legged partridge *A. rufa* [18], and the presence of Japanese quail alleles in the wild migratory common quail *Coturnix coturnix* reveals hybridization between domestic quail and the wild relative [19]. Additionally, mtDNA and nuclear microsatellite analysis indicate gene flow between Silver Pheasant *Lophura nycthemera* and Kalij Pheasant *L. leucomelanos* [20]. Infertile F1 hybrids between the common Pheasant *Phasianus colchicus* and domestic chicken have also been reported in captivity [21].

**Fig 1:**
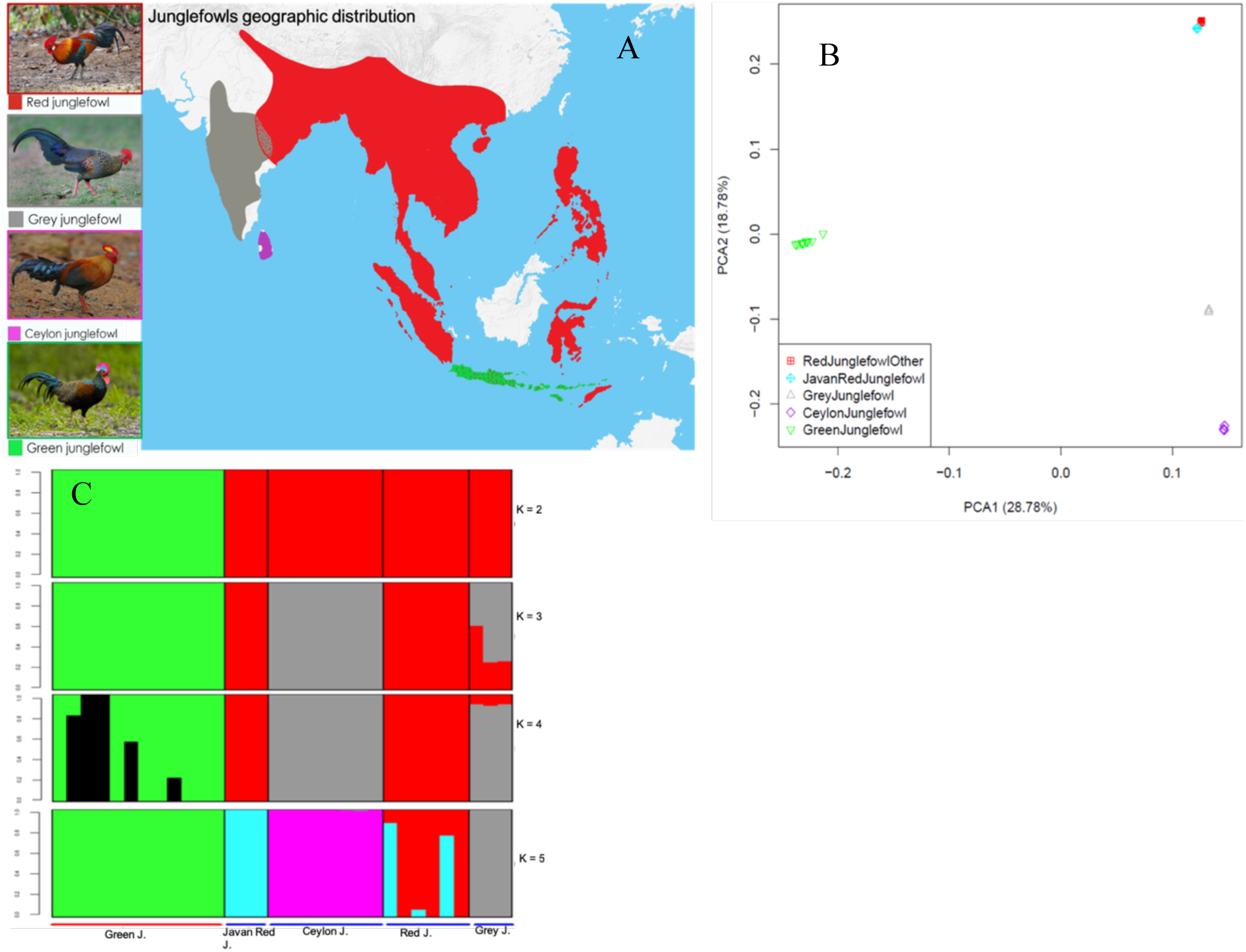
(***A***) The geographic distribution of the four junglefowl species. The sympatric zones where Indian red junglefowl (*Gallus gallus murghi*) overlap with the Grey junglefowl on the Indian subcontinent and Javanese red junglefowl (*Gallus gallus* bankiva) overlap with the Green junglefowl on the Indonesian Islands are annotated with red dots on the map. The map was drawn by overlaying the distribution map of each species obtained from the Handbook of the Birds of the World (consulted in December 2018). Junglefowl species photo credits: Peter Ericsson (Red junglefowl), Clement Francis (Grey junglefowl), Markus Lilje (Ceylon junglefowl), Eric Tan (Green junglefowl). (***B***) Principal component and (***C***) Admixture analysis to establish the species relatedness and genetic structures from the autosomal sequences. 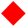: Red junglefowl, 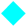: Javanese red junglefowl, 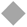: Grey junglefowl, 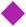: Ceylon junglefowl, 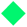: Green junglefowl

Here, we report whole genomes analysis of indigenous domestic village chickens from Ethiopia, Saudi Arabia, and Sri Lanka, together with domestic breeds from Indonesia and China, European fancy chickens and the four wild junglefowl species to infer the genetic contributions of different *Gallus sp.* to the domestic chicken genome. Our results show for the first time the presence of introgressed alleles in domestic chicken from the three non-red junglefowls species (Grey, Ceylon and Green). We also observed extensive introgression from domestic chicken/Red junglefowl into Grey junglefowl, some introgression from domestic chicken into Ceylon junglefowl but no introgression from domestic chicken to Green junglefowl. While our findings support the Red junglefowl as the primary ancestor of domestic chicken worldwide, they also indicate that the genome diversity of domestic chicken population was subsequently reshaped and enhanced following introgression from other *Gallus* species.

## Results

### Sampling, genetic structure and diversity

We analysed 87 whole genome sequences from domestic chickens (n = 53), Red junglefowls (Red (n = 6) and Javanese red (n = 3)), Grey junglefowl (n = 3), Ceylon junglefowl (n = 8), and Green junglefowl (n = 12)) and common Pheasant (n = 2)). Our dataset was made up of newly-sequenced genomes at an average depth of 30X, together with publicly available sequence data, which ranged from 8X to 14X. Across all the 87 genomes, a total of 91,053,192 autosomal single nucleotide polymorphisms (SNPs) were called with more than 50% of the polymorphisms found in common Pheasant (Supplementary Table S1). Summary statistics for read mapping and genotyping are provided in Supplementary Table S1.

To understand the genetic structure and diversity of the four *Gallus* species, we ran principal component (PC) and admixture analyses based on the autosomal SNPs filtered from linkage disequilibrium. PC1 clearly separated the Green junglefowl from the other *Gallus sp*, while PC2 distinguished Red, Grey and Ceylon junglefowls as well as slightly the Javanese red junglefowl subspecies from the other Red junglefowl (Fig. 1*B*), with the Grey and Ceylon junglefowls being positioned closer to each other than to any other junglefowls. The admixture analysis recapitulates these findings, providing some evidence for shared ancestry between the Red and Grey junglefowls at *K* = 3, but at the optimal *K* = 5 the ancestry of each junglefowl species is distinct (Fig. 1*C*).

### Detecting the true *Gallus* species phylogeny

We constructed a neighbour-joining tree and a NeighorNet network using autosomal sequences of 860,377 SNPs filtered to sites separated by at least 1 kb from the total 91 Million SNPs and a maximum likelihood tree on 1,849,580 exon SNPs extracted from the entire autosomal whole-genome SNPs. The trees were rooted with the common Pheasant as an outgroup (Fig. 2*A* and 2*B*; Supplementary Fig. S1*A*). Our results show that Grey and Ceylon junglefowls are sister species and form a clade that is sister to the clade of Javanese red junglefowl, Red junglefowl, and domestic chicken, with the latter two being paraphyletic. Green junglefowl is outside of this clade, making it the most divergent junglefowl species. We also observe the same relationships for the Z chromosome as well as for the mitochondrial (mt) genome (Fig. 2*C* and 2*D*). However, the latter show that the Grey junglefowl samples in this study do carry a domestic/red junglefowl mitochondrial haplotype. All the trees show the Javanese red junglefowl lineage at the base of the domestic/red junglefowl lineages.

**Fig 2.**
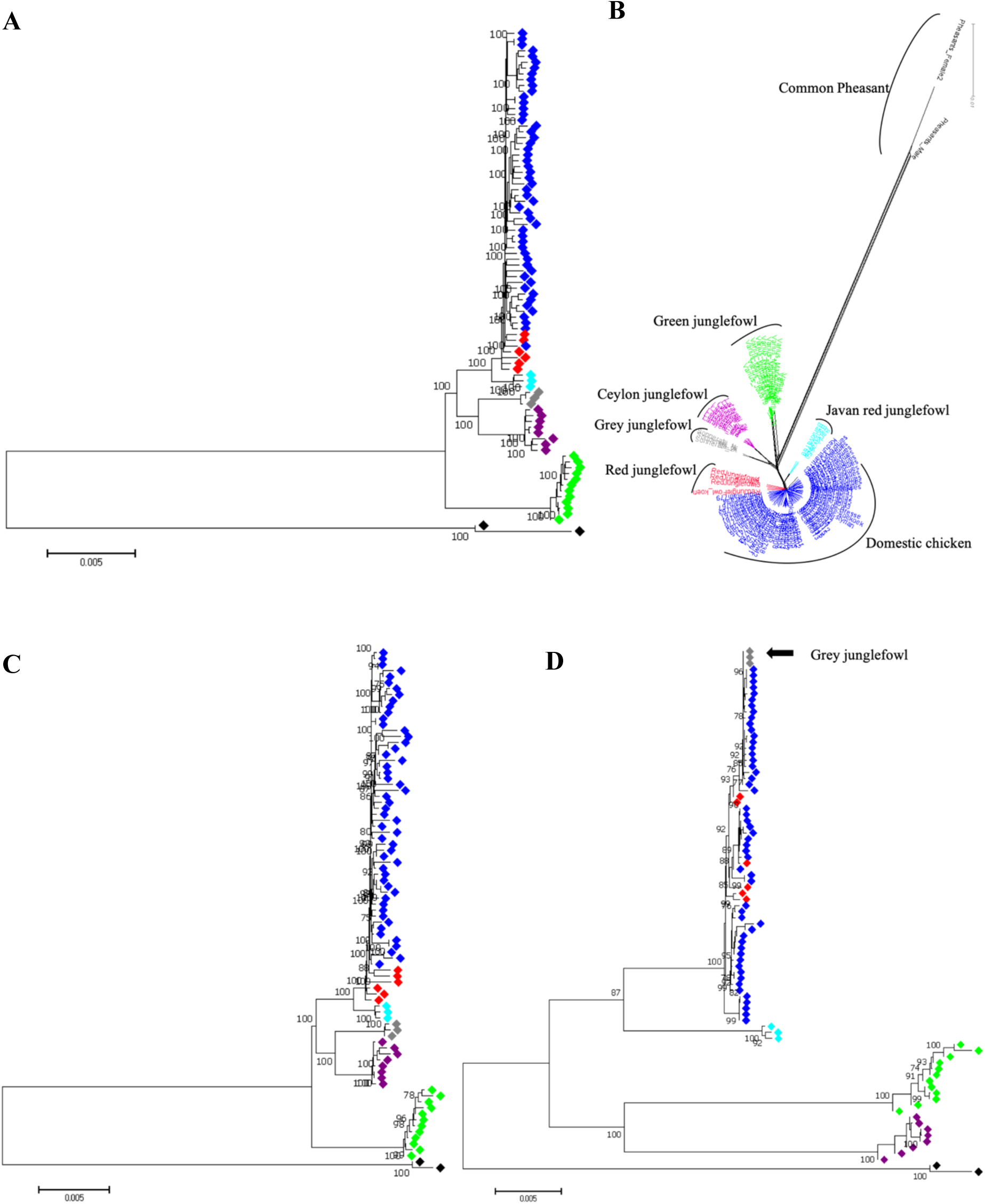
The genome-wide phylogeny of the genus *Gallus.* The figures (***A***), (***C***) and (***D***) are based on Neighbour-Joining phylogenetic trees on the autosomes, Z chromosome and mitochondrial DNA, respectively. The figure (***B***) is the distance matrix of the autosomes constructed from the NeighbourNet network of SplitsTree4. The three Grey junglefowl mtDNA haplotypes in (***D***) are embedded within the domestic/Red junglefowl lineage, indicated with the black arrow. All the trees were rooted with the common Pheasant *Phasianus colchicus*. The colours defining each species are; 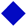: Domestic chicken, 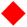: Red junglefowl, 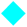: Javanese red junglefowl, 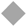: Grey junglefowl, 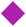: Ceylon junglefowl, 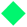: Green junglefowl, 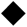: common Pheasant.

Next, we investigated the extent to which other topologies are represented in the autosomal genome using topology weighting by iterative sampling of sub-trees (*Twisst*) [22] based on windows of 50 fixed SNPs [22]. To limit the number of topologies, the analysis was performed twice using either the Red junglefowl or the Javanese red junglefowl along with the Grey, Ceylon and Green junglefowls and the common Pheasant (outgroup). *Twisst* estimates the relative frequency of occurrence (i.e. the weighting) of each of the 15 possible topologies for these five taxa for each window and across the genome.

The most highly weighted topology genome-wide (T12), accounting for ∼ 20% of the genome, supports the autosomal species genome phylogeny: (((((Red or Javanese red junglefowl), (Grey junglefowl, Ceylon junglefowl)), Green junglefowl), common Pheasant) (Fig. 3), while the second highest topology (T9, ∼ 18%) rather places the green junglefowl at the basal of the monophyly Grey and Ceylon junglefowls: ((((Grey junglefowl, Ceylon junglefowl), Green junglefowl), Red or Javanese red junglefowl), common Pheasant). There are also weightings for other topologies. In particular, topologies 3 (2.9%), 10 (7.7%) and 15 (4.2%) show sister relationships between the Red junglefowl and the Grey junglefowl; topologies 6 (2.2%) and 11 (6%) show sister relationships between the Ceylon junglefowl and the Red junglefowl and topologies 1 (3.2%), 4 (3.1%) and 13 (9.7%) show sister relationships between the Green junglefowl and the Red junglefowl.

**Fig 3.**
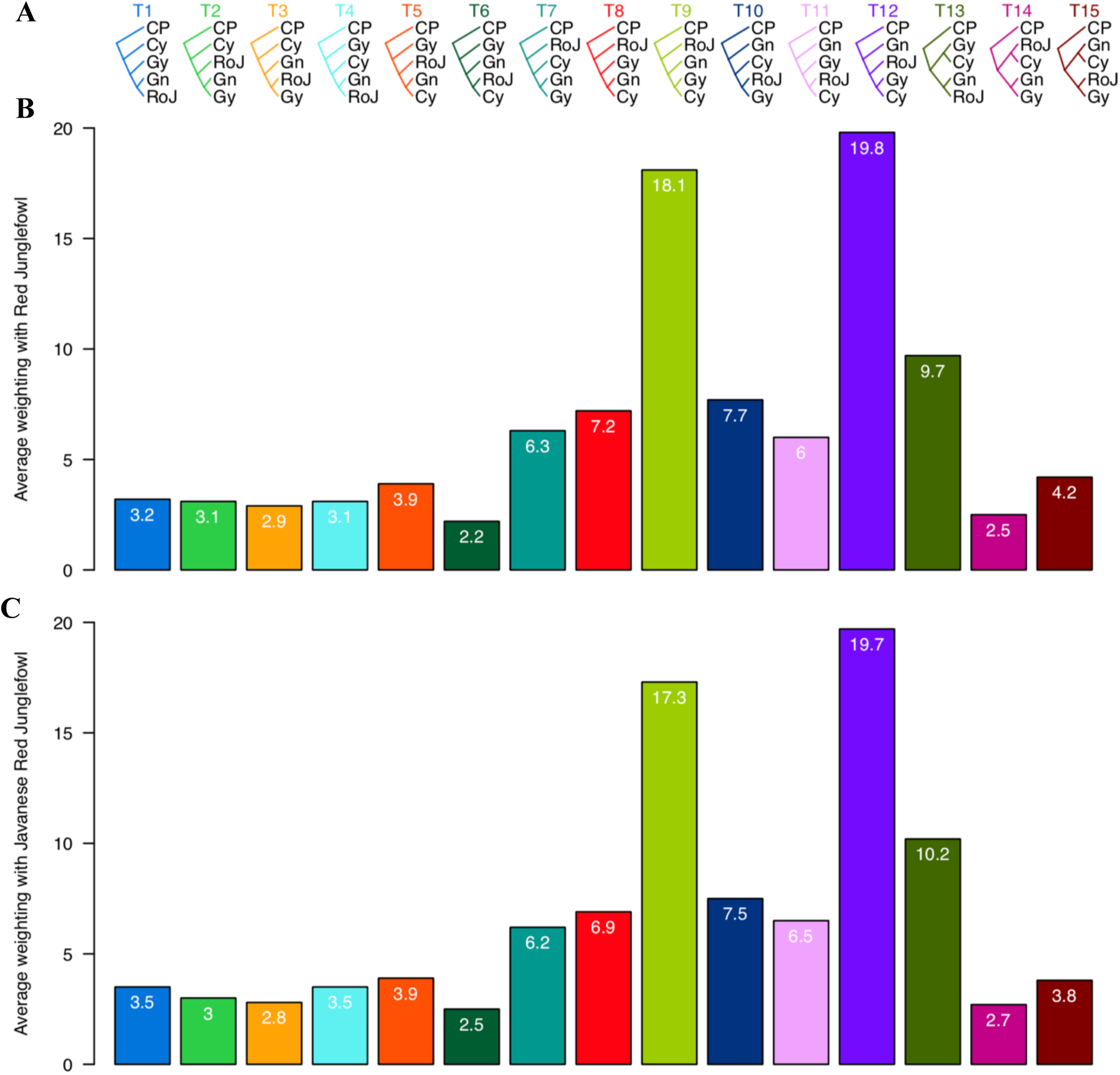
Topology weighting by iterative sampling of sub-trees (*Twisst*) for all the (***A***) 15 possible topologies (T1 – T15) from five taxa: Red junglefowl ‘or’ Javanese red junglefowl (RoJ), Grey junglefowl (Gy), Ceylon junglefowl (Cy), Green junglefowl (Gn) and Common Pheasant (CP). As the number of possible topologies work best for maximum of five taxa (Martin & Van Belleghem, 2017) and with the presence of six taxa in this study, we ran the analysis twice; one with (***B***) Red junglefowl (R) ‘or’ with (**C**) Javanese Red junglefowl (J). (see materials and methods for detail on how this was done). The average weighting (%) for each of the 15 topologies is included in each bar and as well indicated on the *Y* axis.

The result of TreeMix shows similar trends in phylogenetic relationships (as above) but it indicates multiple histories of admixture, namely from Red junglefowl to Grey junglefowl, Ceylon junglefowl to Red junglefowl, and from the root of the monophyly Grey and Ceylon junglefowls to Green junglefowl (Supplementary Fig. S1*B*), with the latter being consistent with the observation on topology 9 in Fig. 3*A*.

### Species divergence time from autosomal genome

To estimate the divergence time between lineages, we first inferred the average coalescence times (CT) from the autosomes, which represents the sum of the accumulated divergence since the split and the standing divergence among the average pair of individuals that was present in the ancestral population at the time of the split. To estimate the split times, we subtracted the estimated nucleotide diversity in each ancestral population from the CT (see materials and methods for details). Among the junglefowls, the divergence times span a few million years. Namely, ∼ 1.1 MYA (Million Years Ago), between the Red and Javanese red junglefowls, ∼ 1.7 MYA between the Ceylon and Grey junglefowls, 2.6 to 2.8 MYA between the Red/Javanese Red and Grey/Ceylon junglefowls, ∼ 4 MYA between Green and the other junglefowls species, while the junglefowls and common Pheasant lineages diverge ∼ 21 MYA (see Table 1 for details of all the pairwise divergence calculations). These split times agree with autosomal and Z chromosome species trees relationships (Fig. 2). Using the same approach, we estimate 8,093 (CI: 7,014 - 8,768) years for the accumulated divergence time (domestication) between the domestic chicken and Red junglefowl (Table 1).

**Table 1.** Divergence time estimates between junglefowl species and with the Common Pheasant. Time in “years”. CT is the coalescence time (i.e. sum of time before and after split), DT is the divergence time (i.e. time from the split to the present).

| Pairwise species comparison | CT (years) | 95% confidence interval (years) | DT (years) | 95% confidence interval (years) |
| --- | --- | --- | --- | --- |
| Domestic chicken - Red junglefowl | 2,060,419 | $1,785,697 \geq CT \leq 2,232,121$ | 8,093 | $7,014 \geq DT \leq 8,768$ |
| Red junglefowl - Javanese red junglefowl | 2,679,697 | $2,322,404 \geq CT \leq 2,903,005$ | 1,164,612 | $1,009,331 \geq DT \leq 1,261,663$ |
| Red junglefowl - Grey junglefowl | 4,072,106 | $3,529,159 \geq CT \leq 4,411,448$ | 2,557,021 | $2,216,085 \geq DT \leq 2,770,106$ |
| Javanese red junglefowl - Grey junglefowl | 4,161,441 | $3,606,582 \geq CT \leq 4,508,228$ | 2,646,356 | $2,293,509 \geq DT \leq 2,866,886$ |
| Grey junglefowl - Ceylon junglefowl | 3,282,030 | $2,844,426 \geq CT \leq 3,555,533$ | 1,766,945 | $1,531,352 \geq DT \leq 1,914,191$ |
| Red junglefowl - Ceylon junglefowl | 4,357,225 | $3,776,262 \geq CT \leq 4,720,327$ | 2,842,140 | $2,463,188 \geq DT \leq 3,078,985$ |
| Javanese red junglefowl - Ceylon junglefowl | 4,379,681 | $3,795,723 \geq CT \leq 4,744,654$ | 2,864,596 | $2,482,650 \geq DT \leq 3,103,312$ |
| Red junglefowl - Green junglefowl | 5,211,015 | $4,516,213 \geq CT \leq 5,645,266$ | 4,057,810 | $3,516,769 \geq DT \leq 4,395,961$ |
| Javanese red junglefowl - Green junglefowl | 5,212,814 | $4,517,772 \geq CT \leq 5,647,215$ | 4,059,609 | $3,518,328 \geq DT \leq 4,397,910$ |
| Grey junglefowl - Green junglefowl | 5,145,901 | $4,459,781 \geq CT \leq 5,574,726$ | 3,992,696 | $3,460,337 \geq DT \leq 4,325,421$ |
| Ceylon junglefowl - Green junglefowl | 5,150,532 | $4,463,794 \geq CT \leq 5,579,743$ | 3,997,328 | $3,464,351 \geq DT \leq 4,330,438$ |
| Red junglefowl - Common Pheasant | 21,885,883 | $18,967,766 \geq CT \leq 23,709,707$ | 20,736,660 | $17,971,772 \geq DT \leq 22,464,715$ |
| Javanese red junglefowl - Common Pheasant | 22,083,637 | $19,139,152 \geq CT \leq 23,923,940$ | 20,934,414 | $18,143,159 \geq DT \leq 22,678,949$ |
| Grey junglefowl - Common Pheasant | 22,136,134 | $19,184,649 \geq CT \leq 23,980,812$ | 20,986,911 | $18,188,656 \geq DT \leq 22,735,820$ |
| Ceylon junglefowl - Common Pheasant | 22,174,484 | $19,217,886 \geq CT \leq 24,022,357$ | 21,025,261 | $18,221,892 \geq DT \leq 22,777,366$ |
| Green junglefowl - Common Pheasant | 22,510,922 | $19,509,466 \geq CT \leq 24,386,832$ | 21,361,699 | $18,513,472 \geq DT \leq 23,141,840$ |

### Genome-wide tests for introgression between junglefowls and domestic chicken

Having established a general pattern for the evolutionary history and relations of the junglefowls, we next assess the presence of shared alleles between the domestic chicken and the junglefowl species. We used *D*-statistics [23, 24] to test for a genome-wide excess of shared alleles between the domestic chicken and each of non-red junglefowl species, relative to the Red junglefowl. *D* is significantly greater than zero with strong Z-scores in all three cases (Table 2), implying possible introgression between domestic chicken and the Grey, Ceylon and Green junglefowls. However, because the Grey and Ceylon junglefowls are sister species, introgression from just one of these species into domestic chicken could produce significantly positive *D* values in both tests. Accordingly, the estimated admixture proportion (*f*) is similar in both cases, ∼ 12% and ∼ 14% for Grey and Ceylon junglefowls, respectively. The estimated admixture proportions are lower for the Z chromosome, ∼ 6% with the Grey junglefowl and ∼10% for Ceylon junglefowl. The estimated admixture proportions between the domestic chicken and the Green junglefowl are ∼ 9% for the autosomes and ∼ 7% for the Z chromosome.

**Table 2.** Patterson’s D statistics and quantification of admixture proportion

| Domestic | Junglefowls | Patterson's *D*-statistics |  |  | Admixture proportion (*f*) |  |
| --- | --- | --- | --- | --- | --- | --- |
|  |  | *D* | Jackknife SD | Z score | *f* estimates | 95% confidence interval |
| Autosomes (chromosomes 1 – 28) |  |  |  |  |  |  |
| Domestic | Grey junglefowl | 0.069 | 0.056 | 37.854 | 0.124 | $0.107 \geq f \leq 0.141$ |
| Domestic | Ceylon junglefowl | 0.055 | 0.046 | 36.776 | 0.142 | $0.134 \geq f \leq 0.150$ |
| Domestic | Green junglefowl | 0.052 | 0.047 | 34.236 | 0.086 | $0.081 \geq f \leq 0.091$ |
| Z chromosome |  |  |  |  |  |  |
| Domestic | Grey junglefowl | 0.041 | 0.089 | 4.177 | 0.059 | $0.031 \geq f \leq 0.086$ |
| Domestic | Ceylon junglefowl | 0.042 | 0.086 | 4.514 | 0.099 | $0.056 \geq f \leq 0.141$ |
| Domestic | Green junglefowl | 0.041 | 0.088 | 4.246 | 0.071 | $0.038 \geq f \leq 0.103$ |

### Genome scans for introgressed regions

To identify specific loci harbouring introgressed variation, we calculated *f*_*d*_ [25] which estimate local admixture proportion within a defined 100 kb windows size. This window size was chosen because it is much greater than the expected size of tracts of shared ancestry from incomplete lineage sorting (ILS) between these species. Given their estimated divergence time and a recombination rate of 3 × 10^−8^, tracts of shared variation across the species that resulted from ILS would be expected to be very small, on the order of ∼ 8 bp (95% CI: 7 – 10 bp) on average (see methods). Next, we separated the domestic chicken into three groups based on their geographic origin and in relation to the geographic location of the junglefowl species: (*i*) Ethiopian and Saudi Arabian domestic chicken at the West of the Grey and wild Red junglefowl geographic distributions (*ii*) Sri Lankan chicken inhabiting the same island as the Ceylon junglefowl, and (*iii*) South-East and East Asian chickens, which include two breeds (Kedu Hitam and Sumatra) from the Indonesian Islands, a geographic area where the Red and the Green junglefowl are found, and Langshan, a breed sampled in the UK but originally from China.

For introgression between domestic chicken and Grey junglefowl, we first selected the three most extreme *f*_*d*_ peaks that are consistent across all three domestic chicken groups for further investigation (Fig. 4): a 26 Mb region on chromosome 1 at chromosomal position 141287737 - 167334186 bp, a 9 Mb region on chromosome 2 at position 11022874 - 19972089 bp, and a 2.8 Mb region on chromosome 4 at position 76429662 - 79206200 bp (Supplementary Table S2*A*; Supplementary Fig. S2*A* – S4*A*). Both the haplotype tree and network show nesting of some Grey junglefowl haplotypes within the domestic chicken lineage, consistent with introgression from domestic chicken/Red junglefowl into Grey junglefowl (Supplementary Fig. S2 – S4 (*B* – *C*)). A result further supported by *Twisst* which indicates localised reductions in the weighting of the species topology and increases in the weightings for both the topologies (((Grey junglefowl, domestic), Red junglefowl), common Pheasant) and (((Grey junglefowl, Red junglefowl), domestic), common Pheasant) (Supplementary Fig. S2*D* – S4*D*). Furthermore, at the candidate introgressed region, *dxy* and *Fst* are reduced between domestic chicken and Grey junglefowl, but not between domestic chicken and Red junglefowl (Supplementary Fig. S2 – S4 (*E* – *F*)). These large tracts therefore show all the signals expected of recent introgression from domestic chicken/Red junglefowl into the Grey junglefowl.

**Fig 4.**
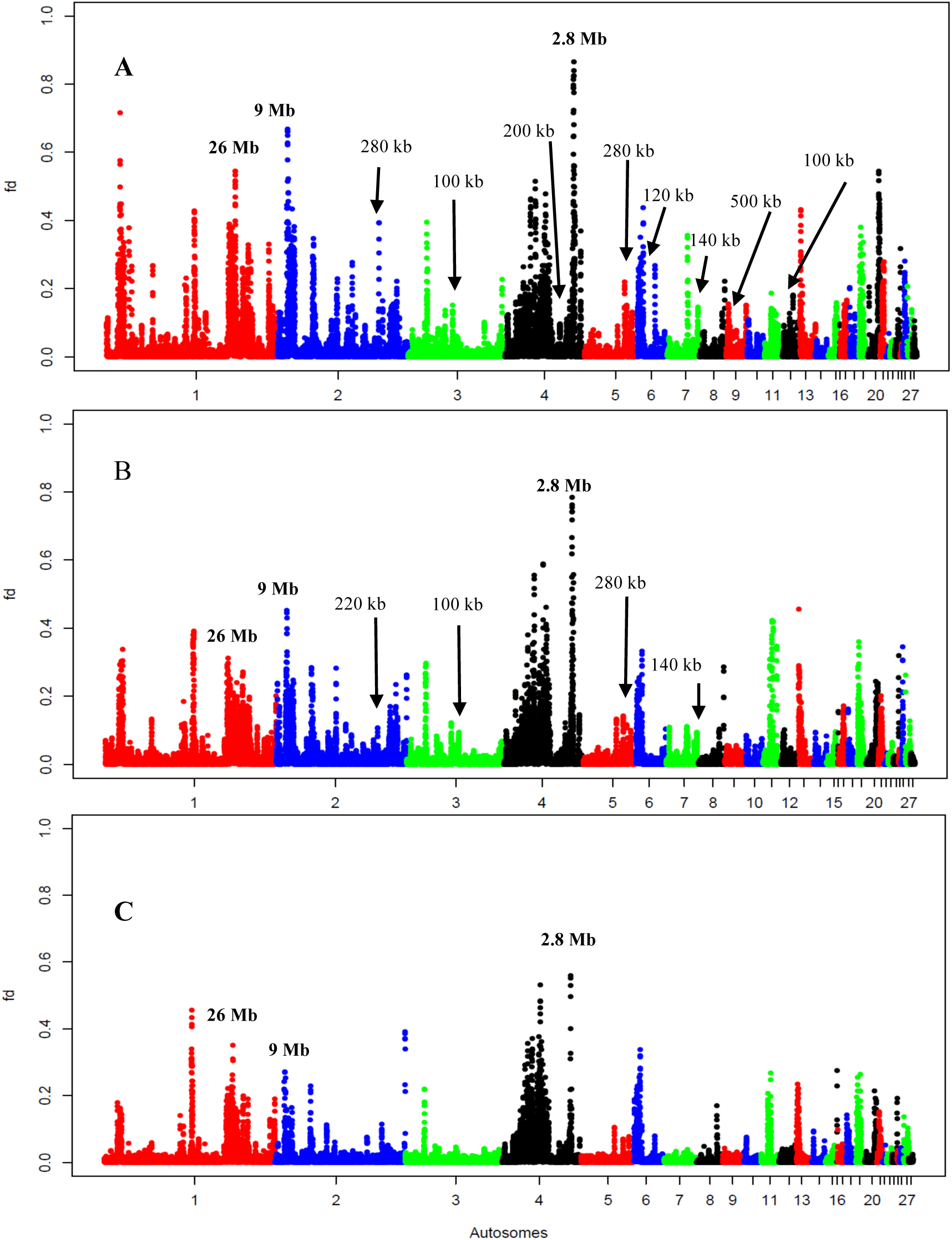
The *f*_*d*_ plots test for the comparison between Grey junglefowl and domestic chicken populations from (***A***) Ethiopia and Saudi Arabia, (***B***) Sri Lanka (***C***) South-East and East Asia. The candidate introgressed regions reported here and their sizes are indicated above each peak (see also Supplementary Table S2). Bold values are introgressed regions from domestic chicken/Red junglefowl into Grey junglefowl, plain values are introgressed regions from Grey junglefowl into domestic chicken. *Y* axis: *f*_*d*_ value spanning 0 to 1, *X* axis: autosomal chromosomes numbers from 1 - 28. See Supplementary figures S12 and S15 for the domestic – Ceylon comparison and domestic – Green junglefowl comparison, respectively.

Next, we investigated candidate introgression signals that are not consistent across the three domestic chicken group comparisons, i.e. peaks present in one or two comparisons but absent in the other(s). Eight candidate regions were randomly analysed and reported here (Supplementary Table S2*B*). These regions are characterised by fragment sizes ranging from 100 kb to 500 kb. Haplotype trees and networks reveal that, unlike the large tracts introgressed into Grey junglefowl, these shorter tracts show some domestic chicken haplotypes (referred to here as targetDom) nested within or close to the Grey junglefowl, indicating introgression from Grey junglefowl into domestic chicken (Fig. 5*A*; Supplementary Fig. S5 – S11). *Twisst* results indicate localised increases in the weighting for the topology (((Grey junglefowl, targetDom), Red Junglefowl), common Pheasant) with proportions ranging from 61% to 80%, much higher than the species topology (((Red junglefowl, targetDom), Grey junglefowl), common Pheasant) ranging from 14% to 28%, and the other alternative topology (((Grey junglefowl, Red junglefowl), targetDom), common Pheasant) ranging from 6% to 11%. These loci are also characterised by reduced *dxy* and *Fst* values between Grey junglefowl and domestic chicken and by increased *dxy* and *Fst* between Red junglefowl and domestic chicken (Fig. 5; Supplementary Fig. S5 – S11 (*E* – *F*)). These Grey junglefowl introgressed haplotypes are most commonly found in Ethiopian chickens, where all eight candidate genomic regions show signal of introgression, followed by Sri Lankan chicken (4 regions), Saudi Arabian chicken (3 regions), Sumatran chicken (2 regions) and chicken from one region in Kedu Hitam and in Red junglefowl (Supplementary Table S2*B*). The introgression found on chromosome 5 was also present in two European fancy chicken breeds (Poulet-de-Bresse and Mechelse-koekoe, Supplementary Fig. S8). No Grey junglefowl introgression was observed in the Langshan chicken. Across these eight regions, a single candidate for bidirectional introgression was observed in the 100 kb region of chromosome 12, where a single Grey junglefowl haplotype is nested within the domestic/Red junglefowl lineage (Supplementary Fig. S11).

**Fig 5.**
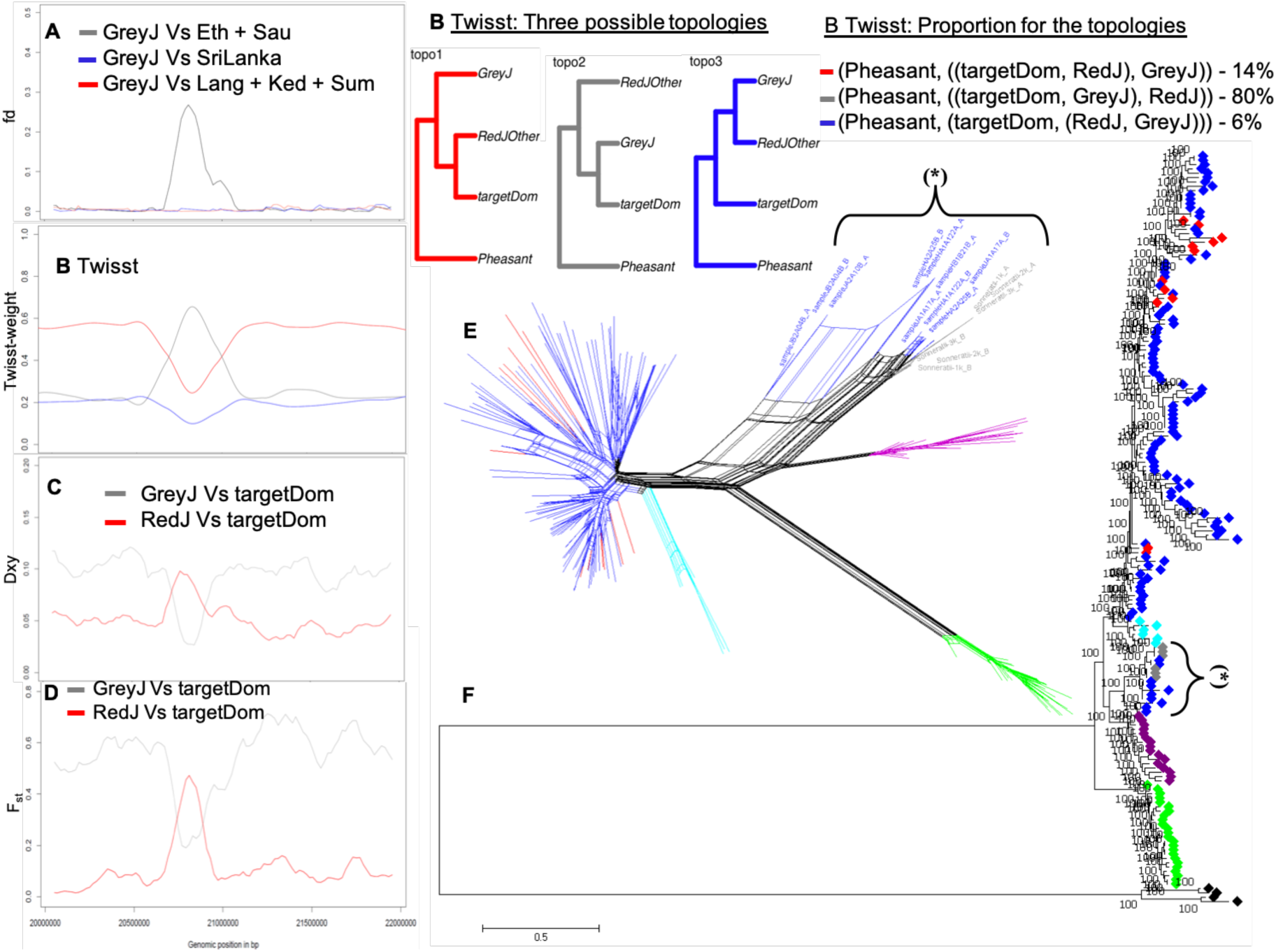
A 120 kb (Chr 6: 21729370 – 21849500 bp) introgressed region from Grey junglefowl into domestic chicken. (***A***) *f*_*d*_ plot for the zoom region, (***B***) *Twisst* plot, its topologies and their proportions. The most consistent topology (80%) has monophyletic relationship between targetDom (introgressed domestic haplotypes) and Grey junglefowl. (***C***) *dxy* and (***D***) *Fst* for the zoom region. Eth, Sau, SriLanka, Lang, Ked and Sum are domestic chickens from Ethiopia, Saudi, Sri Lanka and Langshan, KeduHitam and Sumatra breeds, respectively. targetDom are the introgressed domestic chicken haplotypes from Grey junglefowl (GreyJ) denoted as (**\***) in (***E***) haplotype-based network and (***F***) maximum likelihood tree for the region. The colours defining each population in (***E***) and (***F***) are: 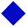: domestic chicken; 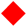: Red junglefowl, 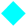: Javanese red junglefowl, 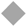: Grey junglefowl, 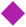: Ceylon junglefowl, 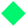: Green junglefowl, 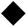: common Pheasant.

A smaller number of candidate regions are detectable in *f*_*d*_ between domestic chicken and Ceylon junglefowl (Supplementary Fig. S12). In most of the candidate regions investigated, haplotype trees and networks indicate unresolved relationships, whereas, some show introgression from Grey rather than Ceylon junglefowl into domestic chicken. By further analysing every peak in the plot, we identified four candidate introgressed regions from Ceylon junglefowl into domestic chicken: three on chromosome 1, spanning 6.52 Mb, 3.95 Mb and 1.38 Mb; and one on chromosome 3, spanning 600 kb (Supplementary Table S2*B*). Both the haplotype trees and networks show introgression of one haplotype into the two different domestic chicken from Sri Lanka for the three candidate regions on chromosome 1 (Supplementary Fig. S13), and two haplotypes into two Sri Lankan domestic chicken for the chromosome 3 region (Fig. 6*B*; Supplementary Fig. S14). The 1.38 Mb region on chromosome 1 also shows introgression from domestic/Red junglefowl into Grey junglefowl (Supplementary Fig. S13*C*). For the four introgressed regions, *Twisst* shows the highest weighting for a topology grouping the target domestic samples with Ceylon junglefowl (Supplementary Fig. S13). Only one candidate region of 100 kb on chromosome 5 shows evidence of introgression from domestic/Red junglefowl into Ceylon junglefowl, supported by both the haplotype network and topology weightings (Fig. 6*C*).

**Fig 6.**
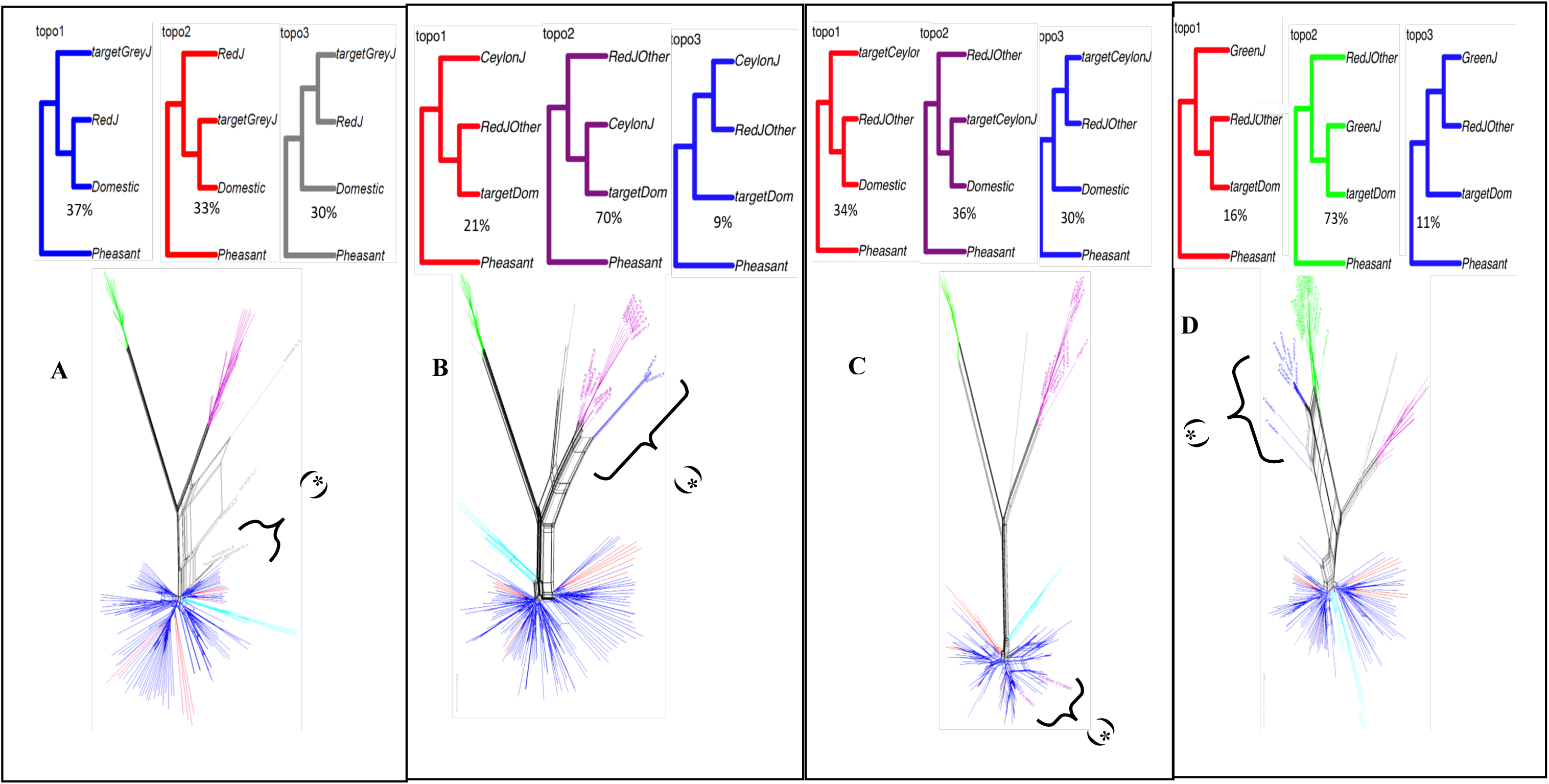
Topologies (*Twisst*), their estimated proportions and network analyses for the introgression from (***A***) domestic chicken to Grey junglefowl (2.8 Mb, Chr 4: 76429662 - 79206200 bp), (***B*)** Ceylon junglefowl to domestic chicken (600 kb, Chr 3: 108325801 - 108925700 bp), (***C***)) domestic chicken/Red junglefowl to Ceylon junglefowl (100 kb, Chr5: 49333700 - 49433700 bp), and (***D***) Green junglefowl to domestic chicken (100 kb, Chr 5: 9538700 - 9638700 bp). (**\***) introgressed haplotypes. The targetGreyJ, targetDom and targetCeylon in the *Twisst* are the introgressed Grey junglefowl, domestic chicken and Ceylon junglefowl haplotypes, respectively as revealed by the network.

There were several peaks of elevated *f*_*d*_ between Green junglefowl and one or more groups of domestic chicken (Supplementary Fig. S15). However, both the haplotype tree and network clearly support introgression only in a single case, from the Green junglefowl into domestic chicken in a 100 kb region on chromosome 5 at position 9538700 - 9638700 bp (Fig. 6*D*). Here, the Green junglefowl is introgressed into 10 of 16 Langshan domestic chicken haplotypes (Supplementary Table S2*B*). As for the candidate regions described above, this case is supported by strong weighting for the topology grouping the target domestic samples with the Green junglefowl, as well as reduced *d*_*XY*_ and *F*_*ST*_ between domestic chicken and Green junglefowl (Supplementary Fig. S16).

### Functional annotations for the enriched genes within the introgressed regions

We identified gene classes that are overrepresented in the regions affected by introgression (Fisher’s exact test, *P* < 0.05). These genes, their gene ontology (GO) terms, functions and their *P*-values are catalogued in Supplementary Table S3. The overrepresented GO terms include some which are linked to the immune responses (e.g. ‘Positive regulation of B cell proliferation’ for the domestic chicken introgression into Grey junglefowl) but also to the regulation of gene expression (e.g. ‘Double-stranded RNA binding’ for the introgression of Grey junglefowl to domestic chicken; ‘Post-embryonic development and Alternative mRNA splicing, *via* spliceosome’ for the introgression of Ceylon junglefowl into domestic chicken).

## Discussion

The Red junglefowl has long been known to be the ancestor of domestic chicken [2-4]. However, one molecular study has shown the presence of an autosomal DNA fragment from the Grey junglefowl in the genome of some domestic chicken [10], and Red junglefowl – domestic chicken mitochondrial DNA have been found in Grey junglefowl [8, 9]. Also, F1 crossbreeding of domestic birds with Green junglefowl is common [5] and captive breeding experiments have reported, although at a very low rate, hatching of eggs and survival of chicks from F1 female Grey x Red junglefowl birds backcrossed to male parental birds from each species [6, 7]. These studies suggest that other species within the genus *Gallus* may have contributed to the diversity of the domestic chicken gene pool. Here, we report for the first time an analysis of the full genomes of the four wild junglefowl species to assess their level of contribution to the domestic chicken.

We first established the species phylogeny of the genus *Gallus.* Both autosomes and Z chromosomes places the Red/Javanese red junglefowl equally close to the Grey and Ceylon junglefowls, which show a sister species relationship. Both also indicate that the Green junglefowl lineage was the first to separate from the common ancestry of the genus. Interestingly, the separation of the Javanese red junglefowl occurs at the root of other Red junglefowl samples studied here, noting that the latter did not include any representative of *G. gallus murghi*, the Red junglefowl subspecies with the largest geographic distribution on the Indian subcontinent. The *Gallus* phylogeny (autosomal) supports a South-East Asian origin of the genus, with a first speciation event separating the Green junglefowl lineage on the present-day Indonesian Islands, occurring ∼ 4 MYA, at the time boundary between the Pliocene and early Pleistocene. Then, a North and North-West dispersion of the continental Red junglefowl ancestral population led to the separation, possibly on the Indian subcontinent, of the lineages leading to the Grey and Ceylon junglefowls ∼ 2.6 to 2.8 MYA. This was followed by the speciation of Grey and Ceylon junglefowls ∼ 1.7 MYA. The South-East Indonesian Islands saw a second arrival of ancestral Red junglefowl ∼ 1.1 MYA, which led to the separation of the Javanese red junglefowl lineage. Using the same approach, we estimated that the domestication of chicken from Red junglefowl likely occurred ∼ 8,000 years ago (95% CI: 7,014 - 8,768 years), earlier than the archaeological evidence (at least 4,000 BP) on the North of the Indian subcontinent [26] but within the Neolithic time and in agreement with the dating of the early chicken remains found from 16 Neolithic sites in China (6,000 BC) [27].

The phylogeny of the genus *Gallus* reported here differ from some other studies [28-30], which are based on short fragments of the genome. In particular, we show here a sister relationship between Grey and Ceylon junglefowls, rather than between Grey and Red junglefowls [28, 30] or between Green and Red junglefowls [29]. A sister relationship between the Grey and Ceylon junglefowls agrees with the today geographic distribution of these two species in South India and Ceylon, respectively. Other studies also indicate more ancient divergence times between the different *Gallus* lineages than the ones reported here (see TimeTree, www.timetree.org). For example, the separation between Grey and Ceylon junglefowl ∼ 1.7 MYA (CI: 1.52 – 1.91 MYA) in this study is more recent than the 8.05 MYA (CI: 3.94 - 12.15 MYA) reported in TimeTree. Several reasons for such discrepancy may be advocated here, e.g. the use of full genome information rather than fragmentary ones as well as different mean Galliformes neutral mutation rates between studies.

Topology weighting analysis, while supporting the same species tree as the primary also show considerable discordance in relationships across the genome, with weightings for topologies that group Red junglefowl/domestic chicken alleles closely with other *Gallus* species. Also, we observe high weighting in *Twisst* analysis for the topology showing relationship between the sister species Grey - Ceylon junglefowls and Green junglefowl, although lower compared to the relation for these two-former species with the Red junglefowl All these results are indicative of presence of incomplete lineage sorting and/or introgression during the history of the genus. Interestingly, while the three non-red junglefowls (i.e. Grey, Ceylon and Green) are allopatric, the fluctuating climatic changes of the Pliocene and early Pleistocene geological era may have not only triggered speciation events within the genus but could have also led to subsequent geographic contact between incipient species, providing opportunities for hybridization at the time. Also, these topologies are supporting introgression resulting from intentional crossbreeding among the genus including between domestic chicken and the Grey, Ceylon and Green junglefowl in modern times.

Several lines of evidence support recent introgression into domestic chicken from other *Gallus* species. Comparison of the *D*-statistic for the autosomes and the Z chromosome show higher levels of admixture on the former than the latter. This trend is not unusual for introgression between species, as species barriers to introgression are often stronger on the sex-chromosomes compared to the autosomes [31].We also report larger genomic tracts showing evidence of introgression than expected under incomplete lineage sorting considering the times for common ancestry reported in this study. This is consistent with recent introgression events where the introgressed haplotypes have not yet been fully broken down by recombination [32]. Typically, haplotypes subject to ILS will be expected to be of smaller size given their antiquity [33]. Obviously, the timeframe of introgression between domestic chicken and the non-red junglefowl species cannot be more ancient than the domestication time and the subsequent dispersion time of domestic birds. At candidate introgressed fragments, we also show excess sequences shared variation between the donors and recipient species, low absolute divergence index with the donor species and genealogical nesting of the candidate introgressed haplotypes within or close to the donor species in both the phylogenies and networks analyses. Together, all the evidence strongly support that these haplotypes represent true introgressed regions from the three non-red junglefowl species into domestic chicken.

Our results also show extensive introgression from domestic chicken/Red junglefowl into Grey junglefowl with introgressed tracts at least as long as 26 Mb in size. It supports recent introgression events into the Grey junglefowl examined here, which originated form captive breed population. The close relationship between domestic chicken and Red junglefowl makes it difficult to pinpoint the source (domestic or red junglefowl) of these introgressed alleles in Grey junglefowl. Specifically, the introgression of Grey junglefowl might have originated in the wild from the Red junglefowl or following the domestication and the dispersion of domestic chicken, considering the long history of sympatry between domestic chicken and the Grey junglefowl across India. Detailed genome analysis of candidate introgressed regions in the wild Grey junglefowl as well as the inclusion in further studies of the Red junglefowl subspecies from the Indian subcontinent *G. g. murghi* may further clarify these issues. Interestingly, amongst the introgressed haplotype regions in the Grey junglefowl, we found several previously proposed chicken domestication genes (e.g. *DACH1, RAB28*) [34, 35] favouring domestic chicken introgression events. Our results highlight the need for further studies of wild Grey junglefowl populations to assess whether their genetic integrity is being threatened by domestic chicken introgression.

We identified introgression from the Grey junglefowl into all domestic chicken populations except in the Langshan, a breed originating from China. It supports the Indian subcontinent as a major centre of origin and dispersion of domestic chicken towards Africa (Ethiopia), the Arabian Peninsula (Saudi Arabia), Sri Lanka, Indonesia and Europe. Interestingly, Ethiopia is the region with the largest proportion of introgressed Grey junglefowl haplotypes in domestic chicken (Supplementary Table S2*B*), possibly a consequence of direct trading routes between the Southern part of the Indian subcontinent and East Africa. It requires further investigation. Surprisingly, we also find evidence of Grey junglefowl introgression into one of the wild Red junglefowl. This Red junglefowl sample originated from the Yunnan Province in China [36], well outside the geographic distribution of the Grey junglefowl confined to India. Here this signature of introgression is likely the result of crossbreeding between domestic chicken and local wild Red junglefowl. Introgression between domestic chicken and wild Red junglefowl has been shown in the past using microsatellite loci in Vietnam [37]. By extension, this result supports a movement of domestic chicken from the centre of origin on the Indian subcontinent towards East and South-East Asia. This hypothesis is also supported by mtDNA analysis which indicates the presence at low frequency of a mtDNA haplogroup in East Asia likely originated from the Indian subcontinent [4].

Our results also highlight the limitations of the current approaches for introgression analysis when dealing with closely related species, the need to include all candidate donor species for the correct interpretation of the introgression patterns, and the importance to complement genome-wide analysis of introgression with locus specific analyses including phylogenetic analysis of haplotypes. The *Gallus* species phylogeny indicates that the Grey and the Ceylon junglefowls are sister species, which speciated after the separation of the Red junglefowl/domestic chicken lineages. Signatures of shared variation suggest that both species have introgressed domestic chicken. However, detailed analysis of candidate introgressed regions reveal that a majority of the Ceylon junglefowl candidate *f*_*d*_ correspond to introgression events involving the Grey junglefowl. This highlights a limitation of both genome wide *D*-statistics and local admixture proportions estimates when there are multiple closely-related donor species. Only a detailed assessment of all the significant *f*_*d*_ candidates using phylogenetic analyses allowed us to identify regions showing introgression from Ceylon junglefowl into domestic chicken. It should also be noted that the genome-wide estimated admixture proportions observed here between the domestic chicken and the Grey, Ceylon and Green junglefowls are likely underestimation following reference genome bias, with all samples aligned against Red junglefowl and in the case of the Grey junglefowl, the use of introgressed reference samples.

At the scale of individual candidate regions, we also observe a different pattern of introgression for Grey and Ceylon junglefowls. While we identify several strong cases of introgression from Grey junglefowl into domestic chicken, evidence for Ceylon junglefowl introgression are limited to one or two Sri Lankan domestic haplotypes at each introgressed region. Similarly, we only reveal one case of introgression from domestic into wild Ceylon junglefowl, a somewhat surprising result considering the potential for introgression in Sri Lanka and the sister relationship between the Ceylon and Grey junglefowls. While we cannot exclude a sampling artefact, the findings suggest that the effect of introgression from Ceylon junglefowl into domestic chicken is likely limited to Sri Lankan domestic birds only. Ceylon junglefowl produce fertile hybrids in captivity with both the Red and Grey junglefowls [5], and there is also anecdotal evidence of human-mediated crosses between male Ceylon junglefowl and female domestic chicken in Sri Lanka (Pradeepa Silva personal communication) to increase the cockfighting vigour of roosters [9].

Crosses between the Green junglefowl and domestic chicken are common in Indonesia [5] suggesting that introgression may have occurred in either direction between these species. The autosomal estimated admixture proportion (*f*) between the domestic chicken and the Green junglefowl is ∼9%. It is ∼7% for the Z chromosome (Table 2). However, our results support only a single compelling example of introgression from the Green junglefowl into domestic chicken. This signal is limited to the Langshan, a Chinese chicken breed. It may represent a legacy of movement of domestic birds from the Indonesian Islands to the East Asian continent. No candidate introgressed regions were detected in the Indonesian domestic chickens (Kedu Hitam and Sumatra). Introgression between the Green junglefowl and domestic chicken may be impeded by genetic barriers, given the greater time since divergence compared to that between Red junglefowl/domestic chicken and the Grey and Ceylon junglefowls.

There is increasing evidence for “adaptive” cross-species introgression amongst mammalian domesticates [38] as well as in humans [33]. A previous study suggests that the chicken yellow skin phenotype is the consequence of introgression event(s) from the Grey junglefowl into domestic chicken [10], a phenotype favoured by some chicken breeders and now fixed in several fancy and commercial breeds [10, 35]. Here, besides some traditional breeds (Langshan, Kedu Hitam Sumatra) with fixed morphological phenotypes, we analysed village chicken populations that are typically characterized by a high level of phenotypic diversity (e.g. plumage colour and pattern, morphology). Introgressed regions were not found fixed or approaching fixation in any of the indigenous village chicken populations examined. Gene ontology (GO) analysis reveals several biological functions related to the control of gene expression (see Supplementary Table S3). Undoubtedly, these candidate introgressed regions are contributing to the genome diversity of the domestic chicken and while we have no evidence of positive selection at these introgressed regions [34], other selection pressures (e.g. heterozygote advantage - balancing selection) may possibly be acting. Whether or these introgression events influence the phenotypic diversity in village chickens is unclear, but it is likely that variation in expression during post embryonic development as revealed by GO analysis contributes to the wide variety of morphological phenotypes observed within and across domestic chickens. For example, among several genes within haplotypes introgressed from Grey junglefowl are *NOX3* and *GSC* involved in ear development and biogenesis of otoconia supporting balance and gravity detection [39, 40]. Moreover, *CPEB3,* which is associated with thermoception and enhancing memory [41, 42] could play central roles in adaptation to new environmental challenges. *MME,* which plays a role in stimulating cytokine production [43] and *RAP2B,* which is mainly expressed in the neutrophils for platelet activation and aggregation [44] might also affect the immune system of introgressed chickens. Other genes of interests include *CDC5L* and *FOXP2* introgressed from Ceylon junglefowl. The former is a key mitotic progression regulator involved in DNA damage response [45] and the latter is a gene involved in song learning in birds [46]. *IPO7* which is introgressed from Green junglefowl is involved in the innate immune system [47].

In conclusion, our study reveals a polyphyletic origin of domestic chicken diversity with major contributions from the Red junglefowl, but also introgression from the Grey, Ceylon and Green junglefowls. These findings provide new insights into the domestication and evolutionary history of the species. Considering the present geographic distributions of the non-red junglefowl species and the dispersal history of domestic chickens, it is unsurprising that the level of introgression amongst domestic populations varies from one geographic region to the other as it will likely reflect their genetic histories. Similarly, analysis of more domestic populations on a wider geographic scale may provide us with a detailed geographic map of the presence and frequency of introgressed regions across the domestic chicken distribution. Our results shed new light on the origin of the diversity of our most important agricultural livestock species and illustrates the uniqueness of each local domestic chicken population across the world.

## Materials and Methods

### Sampling and DNA extraction

Details of the 87 samples studied here and their geographic location of sampling distributions are provided in Supplementary Table S1. Blood samples were collected from the wing vein of 27 indigenous village domestic chickens from three countries (i.e. Ethiopia (n = 11), Saudi Arabia (n = 5) and Sri Lanka (n = 11)) [9, 34, 48], eight Chinese Langshan chicken sampled in the United Kingdom, and 11 non-red junglefowl *Gallus* species (i.e. Grey (n = 2), Ceylon (n = 7) and Green (n = 2) junglefowls). Blood samples from five of the Ceylon junglefowls were obtained from the wild in Uva province of Sri Lanka while the remaining two Ceylon junglefowls blood were sampled from Koen Vanmechelen collection. The two common Pheasants, *Phasianus colchicus* were sampled from the wild in the United Kingdom. Genomic DNA was extracted following the standard phenol-chloroform extraction procedure method [49]. For all, genome sequencing was performed on the Illumina HiSeq 2000/2500/X platforms with an average depth of 30X coverage.

This dataset was complemented with genome sequences from two domestic fancy chicken breeds (Poule de Bresse and Mechelse Koekoek), one Mechelse Styrian, a 16^th^ generation crossbred bird from the Cosmopolitan Chicken Research Project (CCRP: https://www.koenvanmechelen.be/) and as well as one Red, Grey, Ceylon and Green junglefowls sequences also from Koen Vanmechelen collection (www.koenvanmechelen.be/). The publicly retrieved genome sequences of 15 Indonesian indigenous chickens (Sumatra, n = 5 and Kedu Hitam, n = 10) [50], three Javanese red junglefowl *G. g. bankiva* and nine Green junglefowls [50], and five Red junglefowls, sampled in Yunnan or Hainan Provinces (People’s Republic of China)[36] were also included in our dataset. Genome sequence depth for these birds ranges from 8X to 14X.

In total, these 87 genomes include 53 domestic chicken, 6 Red junglefowl, 3 Javanese red junglefowl, 3 Grey junglefowl, 8 Ceylon junglefowl, 12 Green junglefowl and 2 common Pheasants. The newly sequence reads of these birds are accessible at https://www.ncbi.nlm.nih.gov/sra/PRJNA432200 or in the NCBI with the accession number PRJNA432200.

### Sequence mapping and variants calling

Raw reads were trimmed of adapter contamination at the sequencing centre (i.e BGI/Edinburgh Genomics) and reads that contained more than 50% low quality bases (quality value ≤ 5) were removed. Reads from all genomes were mapped independently to the *Galgal* 5.0 reference genome [51] using the Burrows-Wheeler Aligner bwa mem option version 0.7.15 [52] and duplicates were marked using Picard tools version 2.9.0 (http://broadinstitute.github.io/picard/). Following the genome analysis toolkit (GATK) version 3.8.0 best practises [53], we performed local realignment around INDELs to minimize the number of mismatching bases across all reads. To apply a base quality score recalibration step to reduce the significance of any sequencing errors, we used a bootstrapping approach across both the wild non-red junglefowls species and common Pheasants that has no known sets of high-quality database SNPs. We applied same approach to the red junglefowl for consistency. To do this, we ran an initial variant calling on individual unrecalibrated BAM files and then extracted the variants with highest confidence based on the following criteria: -- filterexpression “QD < 2.0 ‖ FS > 60.0 ‖ MQ < 40.0”. We then fed this initial high-quality SNPs as input for known set of database SNPs. Finally, we did a real round of SNPs calling on the recalibrated data. We ran these steps in a loop for multiple times until we reach convergence for each sample.

To improve the genotype likelihoods for all samples using standard hard filtering parameters, we followed the multi-sample aggregation approach, which jointly genotypes variants by merging records of all samples using the ‘-ERC GVCF’ mode in ‘HaplotypeCaller’. We first called variants per sample to generate an intermediate genomic (gVCF) file. Joint genotype was performed for each species separately using ‘GenotypeGVCFs’ and then subsequently merged with BCFtools version 1.4 [54]. Variants were called using Hard filtering “--filterExpression “QD < 2.0 ‖ FS > 60.0 ‖ MQ < 40.0 ‖ MQRankSum < −12.5 ‖ ReadPosRankSum < −8.0”. All downstream analyses were restricted to the autosomes, the Z chromosome and the mitochondrial DNA.

The percentage of the mapped reads and read pairs properly mapped to the same chromosome were calculated using SAMtools “flagstat” version 1.4 [54] while the number of SNPs per sample were identified using VCFtools “vcf-stats” version 0.1.14 [55]. Summary statistics for read mapping and genotyping are provided in Supplementary Table S1.

### Population genetic structure

Principal component analysis was performed on the SNPs identified across the autosomes, filtered with “--indep-pairwise 50 10 0.3”, to visualise the genetic structure of the junglefowl species using PLINK version 1.9 (http://pngu.mgh.harvard.edu/purcell/plink/) and was complemented with analysis using ADMIXTURE version 1.3.0 [56], performed unsupervised with default (folds = 5) for cross-validation in 5 runs with different number of clusters (*K*).

### Species tree

To unravel the species tree of the genus, we constructed an autosomal Neigbour-Joining phylogenetic tree using Phyml version 3.0 [57] and network using NeigborNet option of SplitsTree version 4.14.6 (*splitstree.org)*. First, the dataset was filtered to sites separated by at least 1 kb and then converted to a phylip sequence file using scripts from https://github.com/simonhmartin. We also constructed maximum likelihood tree on the exon variants. This was done by first annotating the entire whole-genome vcf file with SnpEff and then extracted different variants effect within the exons using SnpSift [58]. As with the above, all trees including the Z chromosome were based on polymorphic sites but not for the mtDNA (i.e. all consensus sequences were used). All trees were plotted using the General Time Reversible (GTR) model of nucleotides substitution following its prediction by jModeltest 2.1.7 [59] and then viewed in MEGA 7.0 [60].

After phasing all the autosomal SNPs using SHAPEIT[61], we next performed “Topology Weighting by Iterative Sampling of Sub-Trees” (*Twisst*) [22] which summarized the relationships among multiple samples in a tree by providing a weighting for each possible sub-tree topology. Neighbour-joining trees were generated for fixed 50-SNP windows using Phyml 3.0 [57]. Topologies were plotted in R using the package “APE” version 5.1 [62]. We ran the TreeMix [63] in a block size of 1000 SNPs per window after having filtered the vcf file with “maf 0.01” using PLINK version 1.9 (http://pngu.mgh.harvard.edu/purcell/plink/).

### Species divergence time

To estimate the divergence time between the junglefowl species as well as with the common Pheasant, we first estimated the approximate coalescence time, which include the divergence that has accumulated from the present to the period of split and the divergence among the average pair of individuals that was present in the ancestral population at the time of split, using the equation:

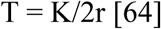

where K is the average sequence divergence for pairwise species. We included both the variant and non-variant sites in the analysis of K which was run in every 100 kb region of the genome with 20 kb step size. r is the Galliformes nucleotide substitution rate per site per year 1.3 (1.2 – 1.5) × 10^−9^ [65], T is the time in year.

Assuming the average diversity (π) across the descendant of junglefowl species is similar to the diversity present in their ancestral population before each split, we estimated the divergence time as below:

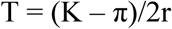

Using the commonest species topology, the average π = (π_Pheasant_ + (π_Green_ + ((π_Grey_ + π_Ceylon_)/2 + (π_Javanese Red_ + π_Red_)/2)/2)/2

### Estimating tract lengths for shared haplotypes under incomplete lineage sorting

Using the approach of Huerta-Sánchez et al [66], we estimated the likely length of shared haplotypes across the genome following incomplete ancestral lineage sorting. This was done with the equation

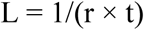

where L is expected length of a shared ancestral sequence, r is recombination rate per generation per bp (3 × 10^−8^ for chicken on the autosomes) [67] and t is the expected divergence time across the junglefowls (∼ 4 MYA), assuming one year generation time.

### Detecting introgression

First, we computed *D*-statistics [23, 24] to test for a genome-wide excess of shared derived allele(s) between two in-groups using the outgroup as representative of the ancestral state. Considering the three in-groups, *P*_*1*_ (Red junglefowl), *P*_*2*_ (domestic chicken) and *P*_*3*_ (Grey or Ceylon or Green junglefowls), and an out-group *O* (common Pheasant), the expected phylogeny is (((*P*_*1*_, *P*_*2*_), *P*_*3*_), *O*). ABBA denotes sites where the derived allele ‘B’ is shared between the domestic chicken ‘*P*_*2*’_ and the Grey or Ceylon or Green junglefowls ‘*P*_*3*’_, while the Red junglefowl ‘*P*_*1*’_ shares the ancestral allele ‘A’ with the common Pheasant ‘*O*’. BABA denotes sites where the Red junglefowl ‘*P*_*1’*_ shares the derived allele ‘B’ with ‘*P*_*3*’_ while the domestic chicken ‘*P*_*2*’_ shares the same ancestral state with the outgroup ‘*O*’. The majority of ABBA and BABA patterns are due to incomplete lineage sorting but an excess of one over the other can be indicative of introgression [23-25]. *D* is the relative excess computed as the difference in the number of ABBA and BABA sites divided by the total number of ABBA and BABA sites. Under the assumption of no gene flow and a neutral coalescent model, count of both ABBA and BABA should be similar and *D* should tend to zero. We used the approach of Durand et .al [24] to compute ABBA and BABA counts from allele frequencies, in which each SNP contributes to the counts even if it is not fixed. We used the jackknife approach, with a block size of 1 Mb to test for a significant deviation of *D* from zero (i.e consistent with introgression), using a minimum Z score of 4 as significant. We then estimated the proportion of admixture, *f* [23, 24]

### Identifying introgression at particular loci and inferring the direction of introgression

To identify specific regions showing introgression between the domestic chicken and the non-red junglefowl species, we used a combination of analyses. First, we estimated *f*_*d*_ [25], which is based on the four-taxon ABBA-BABA statistics and was designed to detect and quantify bidirectional introgression at particular loci [25], *f*_*d*_ was computed in 100 kb windows with a 20 kb step size. Each window was required to contains a minimum of 100 SNPs. The strongest candidate regions of introgression (highest elevated *fd* values) were visually assessed to determine the introgression size. We avoided the use of extreme tail end or cut-off approach to prevent losing out peaks harbouring introgression into domestic chicken as the majority of candidate introgression signals supports introgression from domestic chicken into Grey junglefowl. These *f*_*d*_ regions were then extracted and further investigated using *Twisst* [22] to test for a deviation in topology weightings in the candidate regions. Here, we used only four taxa: domestic chicken, Red junglefowl, common Pheasant, and either the Grey, Ceylon or Green junglefowl.

Next, we constructed haplotype-based gene trees and networks to make inferences about the direction of gene flow. The expectation is that introgressed regions in domestic chicken from any of the non-red junglefowls will be indicated by finding chicken haplotypes nested within the donor species, or with the donor species haplotypes at the root of the introgressed ones. For regions in non-red junglefowls that are introgressed from domestic chicken, the expectation is that the introgressed haplotypes will be nested within the domestic chicken clade. To do this, sequences from the candidate introgressed regions were phased using SHAPEIT [61]. The phased haplotypes were converted into a variant call format file and subsequently formatted in Plink 1.9 [68] with the ‘beagle recode’ option, the output from which was provided as an input to a custom bash script to generate a FASTA file. The optimal molecular evolutionary model was inferred using jModeltest 2.1.7 [59] based on the Akaike information criterion (AIC). Phyml 3.0 [57] was used to compute the approximate likelihood ratio score for each branch using the best predicted model. For the network, we used the NeigborNet option of SplitsTree version 4.14.6 (*splitstree.org)*. The input file for the network was a distance matrix created using ‘distMat.py’ accessible at https://github.com/simonhmartin/genomics_general.

Finally, we examined levels of divergence between species to further validate our candidate regions. Introgression between domestic chicken and either the Grey, Ceylon or Green junglefowls is expected to reduce genetic divergence between the two species, regardless of the direction of introgression. Introgression into domestic chicken is expected to also increase divergence between domestic chicken and Red junglefowl, whereas introgression from domestic chicken into the Grey, Ceylon or Green junglefowl should not affect divergence between domestic chicken and Red junglefowl. We therefore computed relative (*F*_*ST*_) and absolute (*d*_*XY*_) measures of divergence between pairs using the script popgenWindows.py available at https://github.com/simonhmartin/genomics_general.

### Remapping of candidate introgressed regions to GRCg6a

Following the recent release of new reference genome (GRCg6a), all candidate introgressed regions obtained from *Galgal* 5.0 were remapped using the NCBI remapper tool. All remapping options were set to the default threshold. The GRCg6a coordinates for the candidate introgressed regions and genes are reported here throughout the manuscript.

### Gene ontology analysis

All candidate genes within the introgressed regions for different pairwise comparison and in different introgressed directions were used to determine their biological cluster functions using DAVID version 6.8 (https://david.ncifcrf.gov/summary.jsp). Only gene ontology with Fisher exact *P* < 0.05 default threshold were retained in this study.

## Acknowledgements and Funding

This study was conducted during Raman Akinyanju Lawal PhD programme, supported by the University of Nottingham Vice Chancellor’s Scholarship (International) award. Financial support for sampling and/or genome sequencing were obtained from the University of Nottingham, Biotechnology and Biological Sciences Research Council (BBSRC), the UK Department for International Development (DFID) and the Scottish Government (CIDLID program, BB/H009396/1, BB/H009159/1 and BB/H009051/1), BMGF Grant Agreement OPP1127286, the National Plan for Science, Technology and Innovation (MAARIFAH), King Abdulaziz City for Science and Technology, Kingdom of Saudi Arabia.

## Author contributions

O.H. and R.A.L. designed and supervised the project with major contributions from S.H.M for the data analysis. P.S. contributed the DNA of five Ceylon junglefowls and domestic birds from Sri Lanka. R.M.A., R.S.A., and J.M.M. collected the samples and provided the DNA of the Saudi Arabian birds. All the captive junglefowl blood samples were collected from K.V. farm while their DNA preparation was performed by R.A.L., D.W. and O.H. The DNA preparation of the Ethiopian chickens were performed by J.M.M. The genome sequences of the fancy birds, one Red, Grey, Ceylon and Green junglefowls were contributed by K.V. and A.V. Langshan samples were collected by P.M.H with genome sequences information provided by D.D.W. and Y-P.Z. The genome sequence of Pheasant was provided by J.S. All data analyses were performed by R.A.L. The manuscript was prepared by R.A.L and substantially revised by O.H and S.H.M. All other authors reviewed and accepted the final draft of the manuscript.

