## Supplementary Materials for "The wild species genome ancestry of domestic chickens"

**Full title: The wild species genome ancestry of domestic chickens**
**Short title: Chicken genome ancestry**

Raman Akinyanju Lawal[1,2*#], Simon H. Martin[3,4], Koen Vanmechelen[5], Addie Vereijken[6], Pradeepa Silva[7], Raed Mahmod Al-Atiyat[8], Riyadh Salah Aljumaah[9], Joram M. Mwacharo[10], Dong-Dong Wu[11,12], Ya-Ping Zhang[11,12], Paul M. Hocking[13†], Jacqueline Smith[13], David Wragg[14] & Olivier Hanotte[1, 14,15*]

[1]Cells, Organisms and Molecular Genetics, School of Life Sciences, University of Nottingham, NG7 2RD, Nottingham, United Kingdom
[2,#]The Jackson Laboratory, 600 Main Street, Bar Harbor, ME 04609, USA
[3]Institute of Evolutionary Biology, University of Edinburgh, EH9 3FL, Edinburgh, United Kingdom
[4]Department of Zoology, University of Cambridge, CB2 3EJ, Cambridge, United Kingdom
[5]Open University of Diversity - Mouth Foundation, Hasselt, Belgium
[6]Hendrix Genetics, Technology and Service B.V., P.O. Box 114, 5830, AC, Boxmeer, The Netherlands
[7]Department of Animal Sciences, Faculty of Agriculture, University of Peradeniya, Sri Lanka
[8]Genetics and Biotechnology, Animal Science Department, Agriculture Faculty, Mutah University, Karak, Jordan
[9]Department of Animal Production, King Saud University, Saudi Arabia
[10]Small Ruminant Genomics, International Centre for Agricultural Research in the Dry Areas (ICARDA), P.O. Box 5689, ILRI-Ethiopia Campus, Addis Ababa, Ethiopia
[11]Center for Excellence in Animal Evolution and Genetics, Chinese Academy of Sciences, 650223 Kunming, China
[12]State Key Laboratory of Genetic Resources and Evolution, Kunming Institute of Zoology, Chinese Academy of Sciences, 650223 Kunming, China.
[13]The Roslin Institute and Royal (Dick) School of Veterinary Studies, University of Edinburgh, Easter Bush Campus, Midlothian, EH25 9RG, UK
[14]Centre for Tropical Livestock Genetics and Health, The Roslin Institute, EH25 9RG, Edinburgh, UK
[15]LiveGene, International Livestock Research Institute (ILRI), P. O. 5689, Addis Ababa, Ethiopia

[†]Deceased
[#]Present address

**The content of this supplementary materials is as below:**

**Fig. S1A and B**: Exon phylogeny and TreeMix
**Fig. S2 – S4**: Plots for the introgression from domestic chicken/Red junglefowl to Grey junglefowl
**Fig. S5 – S11**: Plots for the introgression from Grey junglefowl to domestic chicken/Red junglefowl
**Fig. S12**: Genome *fd* plots for the comparison between Ceylon junglefowl and domestic chicken
**Fig. S13 – S14**: Plots for the introgression from Ceylon junglefowl to domestic chicken.
**Fig. S15**: Genome *fd* plots for the comparison between Green junglefowl and domestic chicken
**Fig. S16**: Plots for the introgression from Green junglefowl to domestic chicken
**Table S1**: Sampling, mapping and variants statistics
**Table S2A and B**: Candidate introgressed regions
**Table S3:** Functional annotations for the enriched genes within the introgressed regions

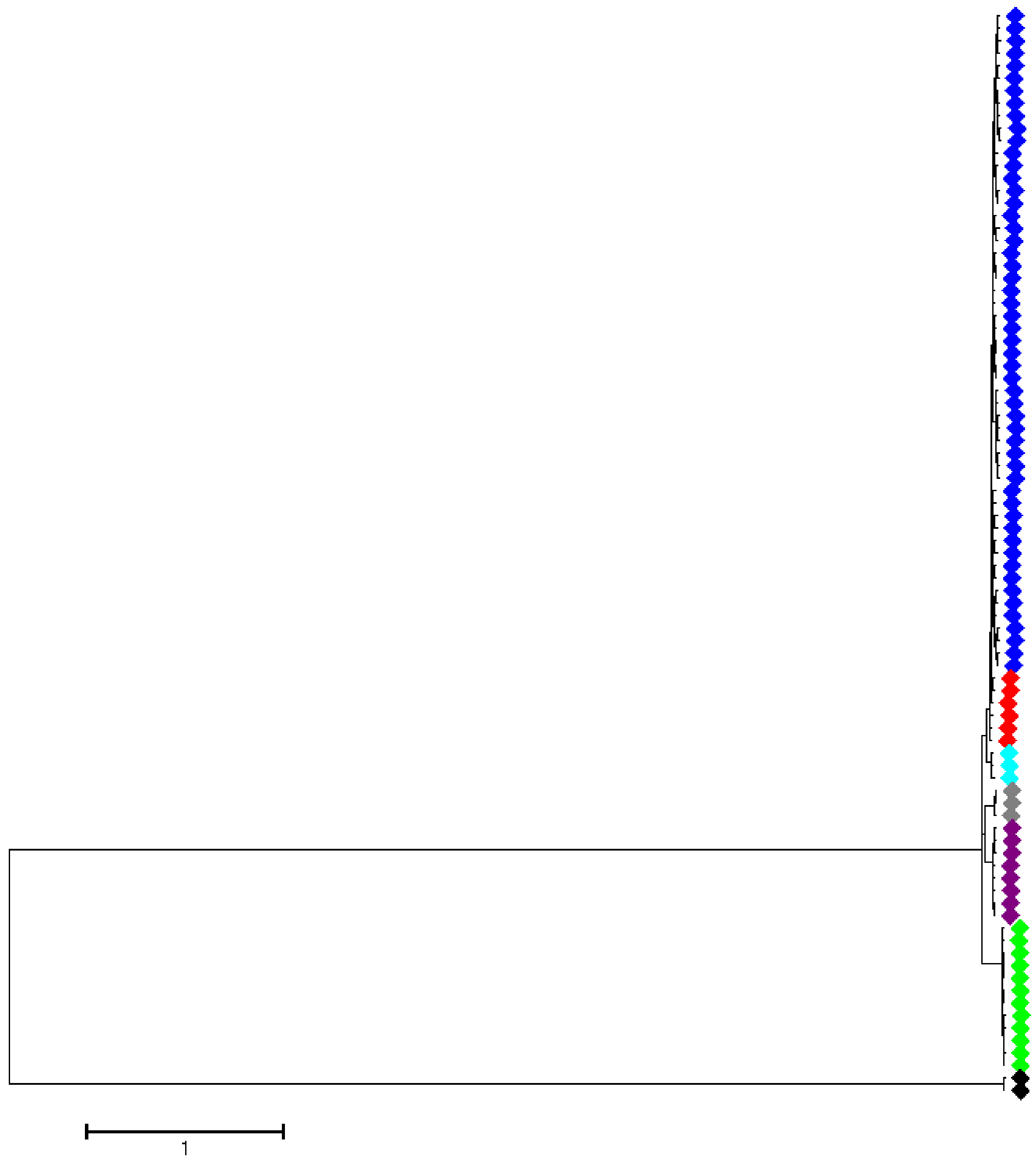


**Fig. S1A.** Maximum likelihood tree generated from 1,849,580 exon SNPs with GTR model. All branches are supported by 100% bootstrap values. The colour for the taxon markers are ◆: Domestic chicken, ◆: Red junglefowl, ◆: Javan red junglefowl, ◆: Grey junglefowl, ◆: Ceylon junglefowl, ◆: Green junglefowl, ◆: common Pheasant.

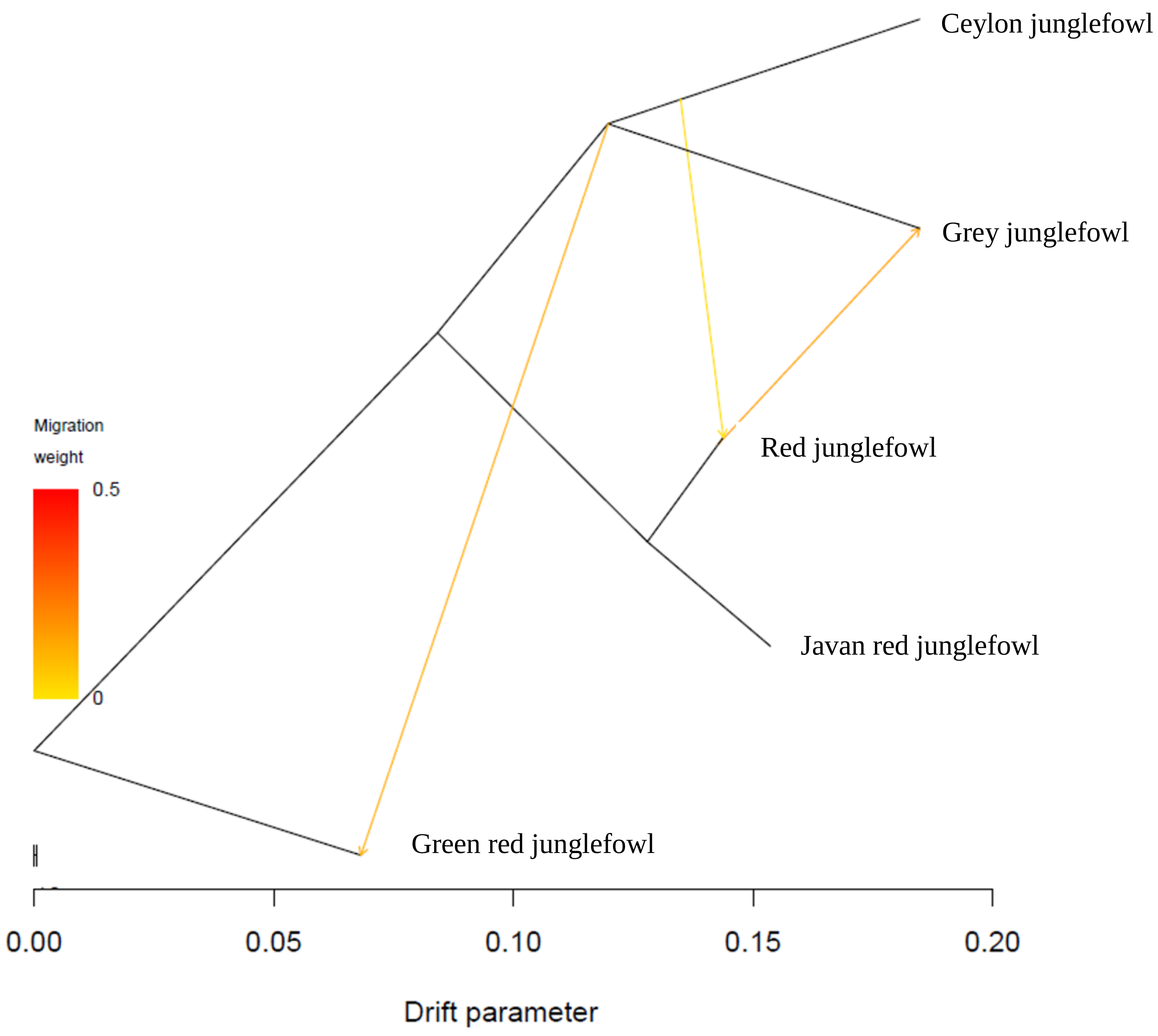


**Fig. S1B.** TreeMix across the autosomal genome

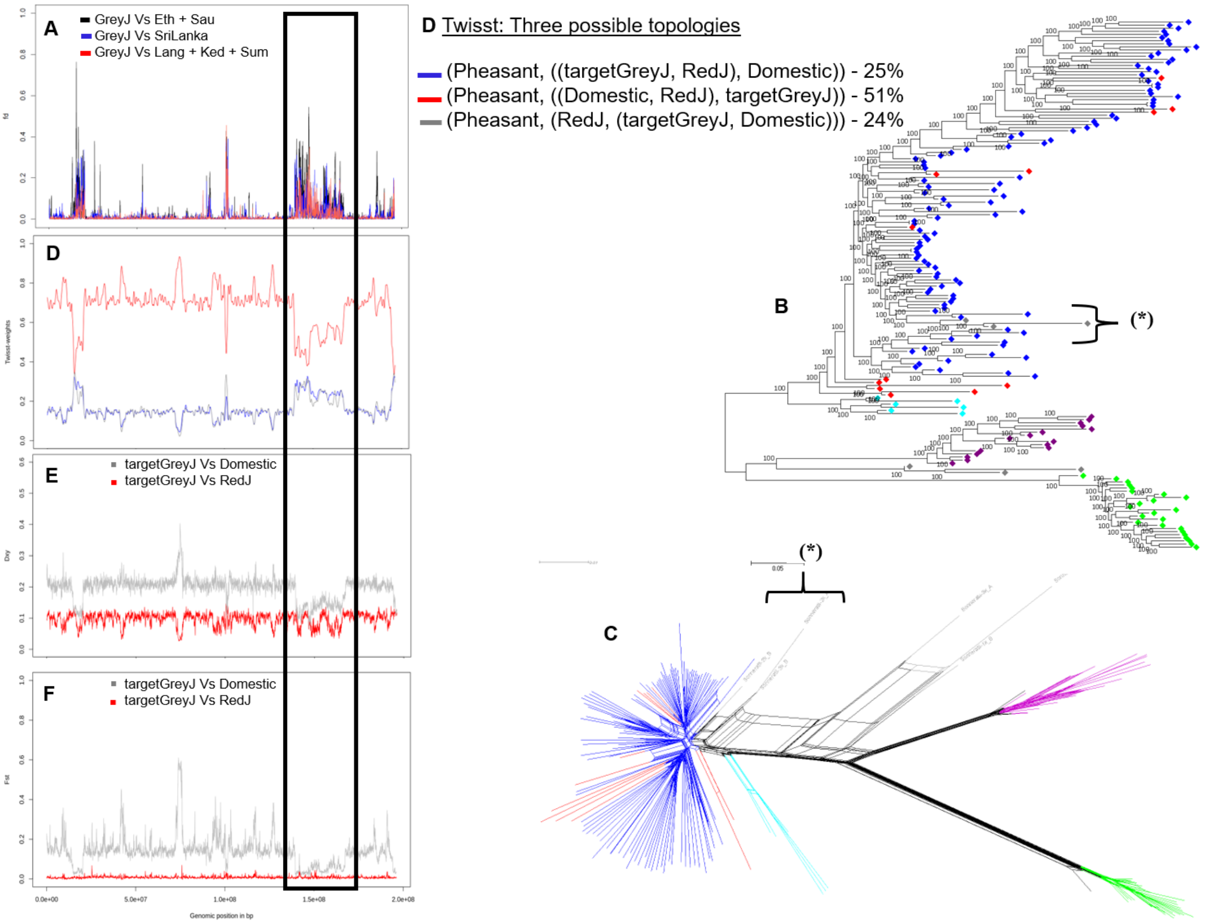


**Fig. S2.** A 26 Mb (Chr1: 141287737 - 167334186 bp) introgressed region from domestic chicken into Grey junglefowl. (**A**) fd plot for chromosome 1, (**B**) haplotype-based network and (**C**) maximum likelihood tree for the introgressed region, (**D**) *Twisst* plot and the proportion of each topology in the introgressed region, (**E**) $D_{XY}$ and (**F**) $F_{ST}$. Eth, Sau, SriLanka, Lang, Ked, Sum represent chicken samples from Ethiopia, Saudi, Sri Lanka, Langshan, Kedu Hitam and Sumatra, respectively, GreyJ represent Grey junglefowl, and targetGreyJ are the introgressed (*****) Grey junglefowl haplotypes. domestic include all the domestic chicken populations. The colours for (**B**) and (**C**) defining each species are; ◆: Domestic chicken ◆: Red junglefowl, ◆: Javan red junglefowl, ◆: Grey junglefowl, ◆: Ceylon junglefowl, ◆: Green junglefowl, ◆: common Pheasant.

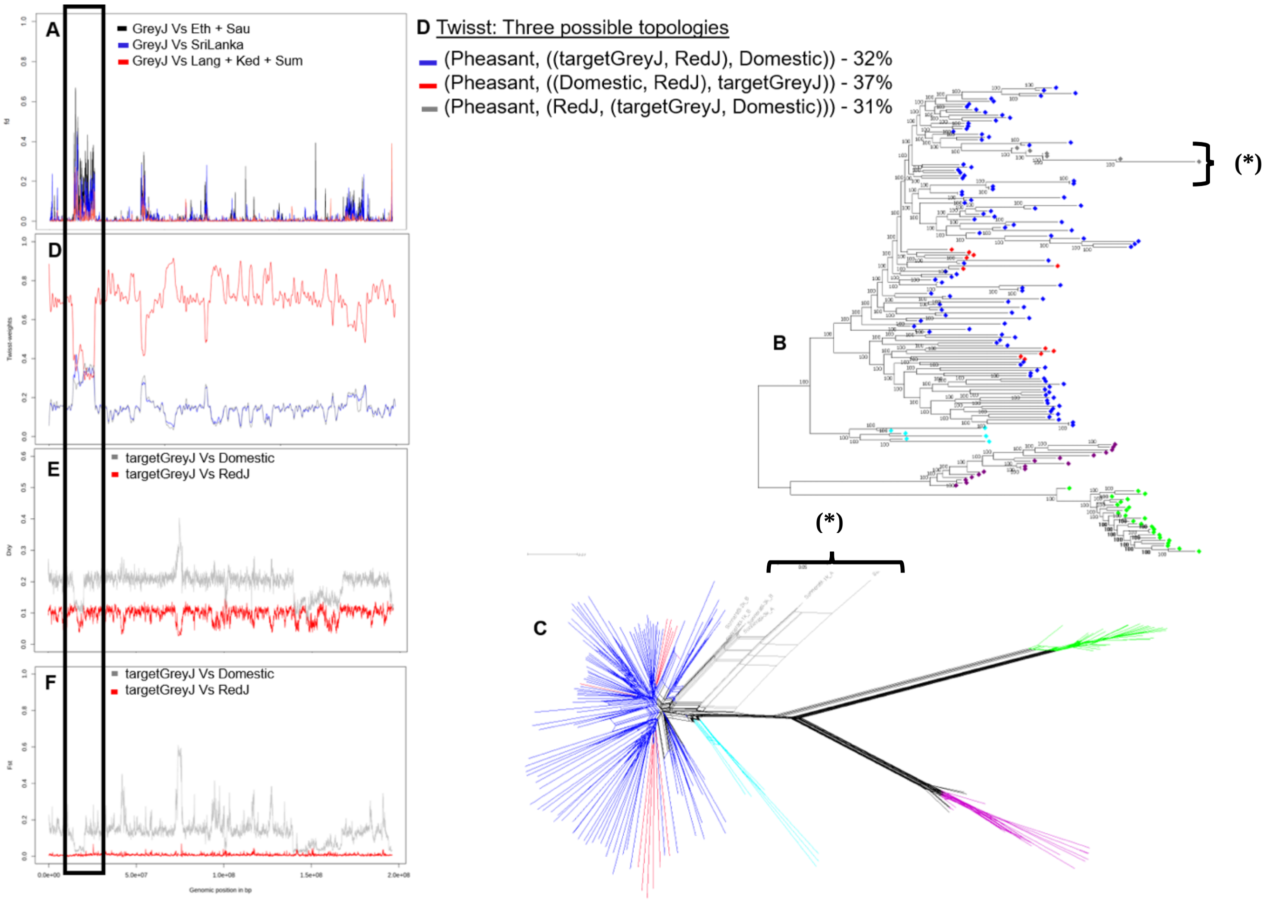


**Fig. S3.** A 9 Mb (Chr2: 11022874 - 19972089 bp) introgressed region from domestic chicken to Grey junglefowl. (**A**) fd plot for chromosome 1, (**B**) maximum likelihood tree for the introgressed region, (**C**) haplotype-based network, (**D**) *Twisst* plot and the proportion of each topology in the introgressed region, (**E**) $D_{XY}$ and (**F**) $F_{ST}$. Eth, Sau, SriLanka, Lang, Ked, Sum represent chicken samples from Ethiopia, Saudi, Sri Lanka, Langshan, Kedu Hitam and Sumatra, respectively, GreyJ represent Grey junglefowl, and targetGreyJ are the introgressed (*****) Grey junglefowl haplotypes. Domestic include all the domestic chicken populations. The colours for (**B**) and (**C**) defining each species are; ◆: Domestic chicken ◆: Red junglefowl, ◆: Javan red junglefowl, ◆: Grey junglefowl, ◆: Ceylon junglefowl, ◆: Green junglefowl, ◆: common Pheasant.

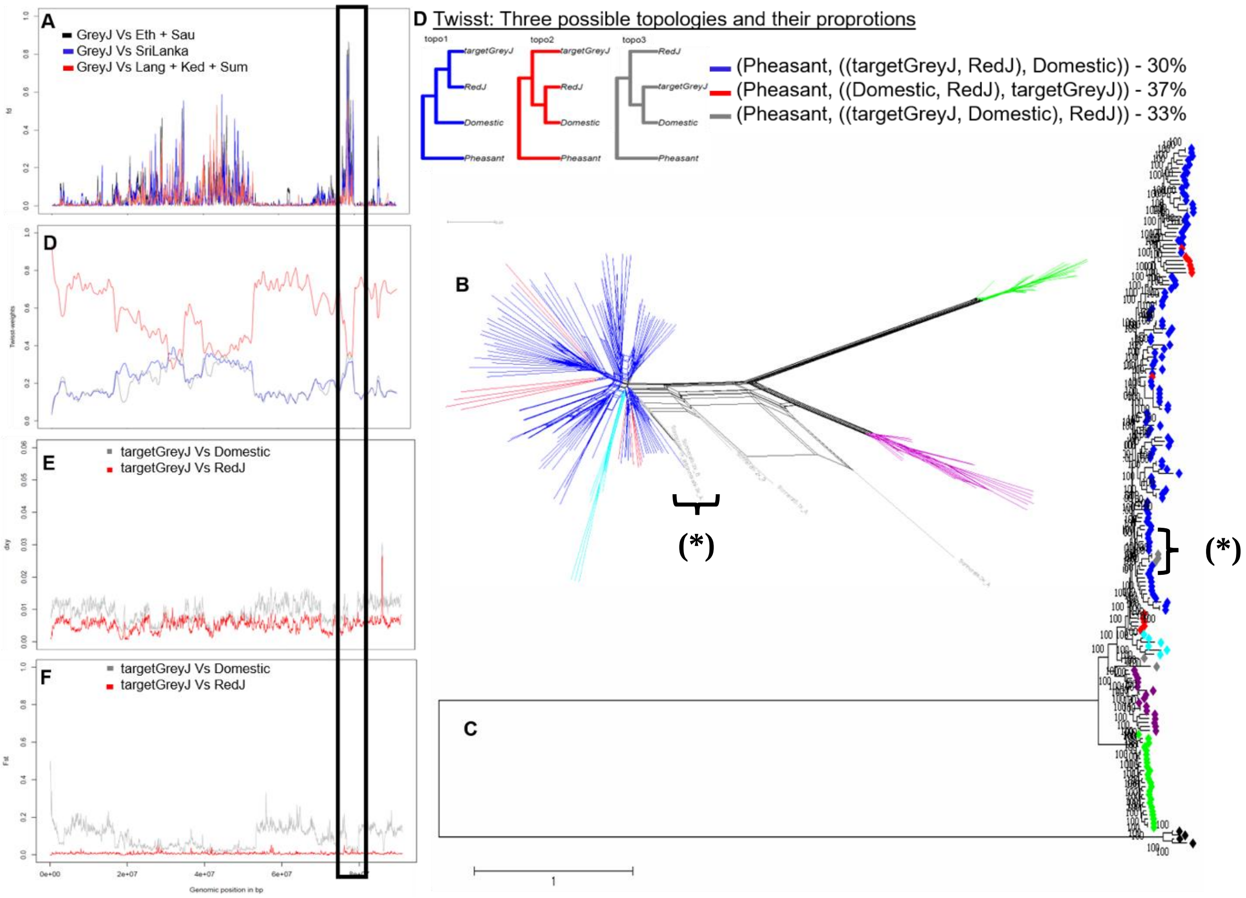


**Fig. S4.** A 2.8 Mb (Chr4: 76429662 - 79206239bp) introgressed region from domestic chicken/Red junglefowl into Grey junglefowl. (**A**) $f_d$ plot for chromosome 4, (**B**) haplotype-based network, (**C**) maximum likelihood tree, (**D**) *Twisst* plot, its topologies and their proportions, (**E**) $D_{XY}$ and (**E**) $F_{ST}$. Eth, Sau, SriLanka, Lang, Ked and Sum are domestic chicken from Ethiopia, Saudi, Sri Lanka and Langshan, KeduHitam and Sumatra breeds, respectively. GreyJ is Grey junglefowl, and targetGreyJ are the introgressed Grey junglefowl haplotypes (*****). Domestic includes all the chicken domestic populations. ◆: Domestic chicken; ◆: Red junglefowl, ◆: Javan red junglefowl, ◆: Grey junglefowl, ◆: Ceylon junglefowl, ◆: Green junglefowl, ◆: common Pheasant.

Description for **Fig. S5 - S11**

The plots are zoomed close to the region. (**A**) fd plot, (**B**) haplotype-based network and (**C**) maximum likelihood tree for the introgressed region (**D**) *Twisst* plot and the proportion of each topology in the introgressed region (**E**) $D_{XY}$ and (**F**) $F_{ST}$. Eth, Sau, SriLanka, Lang, Ked, Sum represent chicken samples from Ethiopia, Saudi, Sri Lanka, Langshan, Kedu Hitam and Sumatra, respectively, GreyJ represent Grey junglefowl, and targetDom are the introgressed (*****) domestic haplotypes. The colours for (**B**) and (**C**) defining each species are ◆: Domestic chicken ◆: Red junglefowl, ◆: Javan red junglefowl, ◆: Grey junglefowl, ◆: Ceylon junglefowl, ◆: Green junglefowl, ◆: common Pheasant.

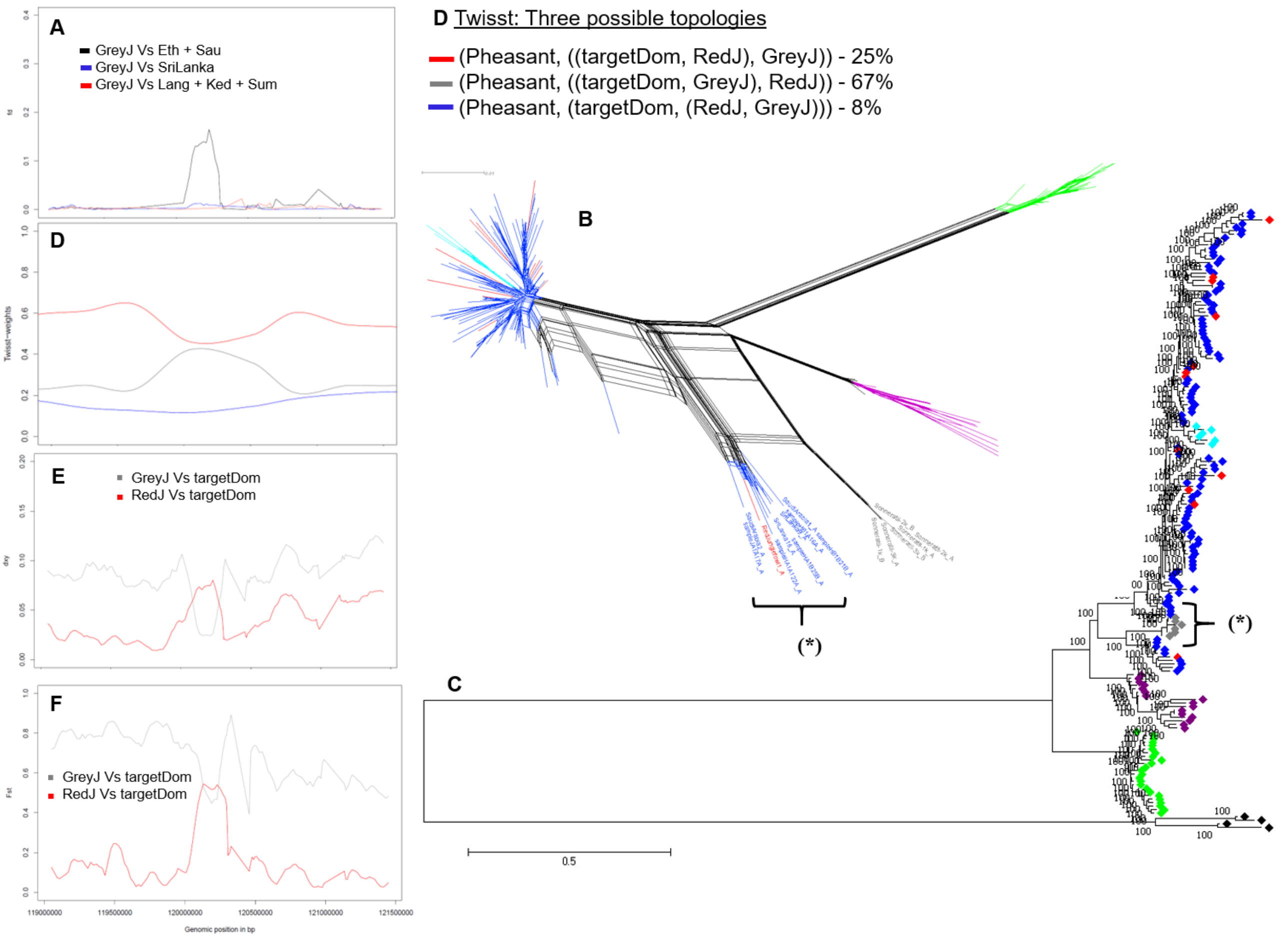


**Fig. S5.** A 220 kb (Chr 2: 119676880 - 119901132bp) candidate introgressed region from Grey junglefowl to domestic chicken/Red junglefowl. targetDom here include the introgressed domestic chicken and a single Red junglefowl haplotypes (*****).

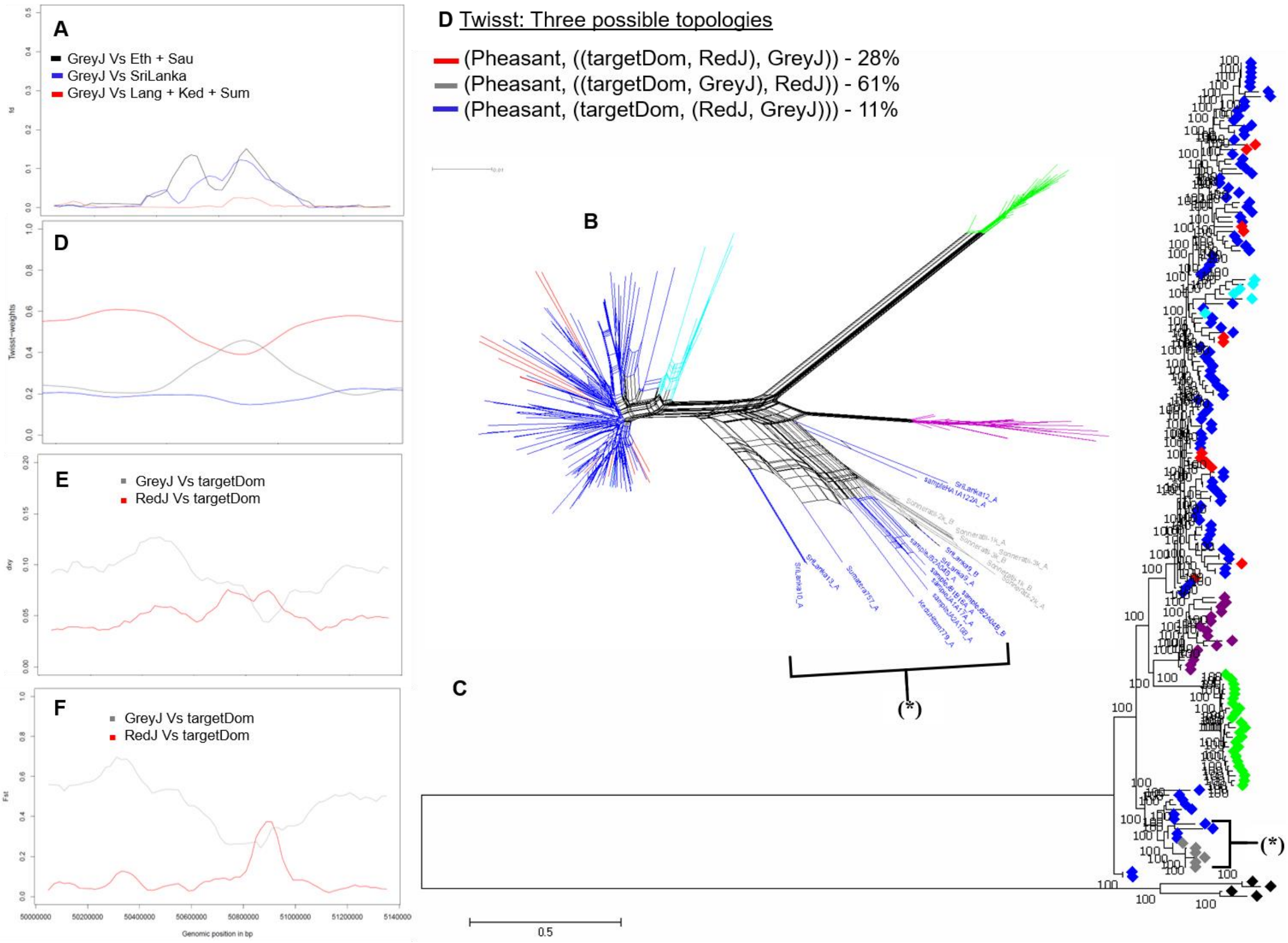


**Fig. S6.** A 100 kb (Chr 3: 50759656 - 50859645 bp) introgressed region from Grey junglefowl into domestic chicken.

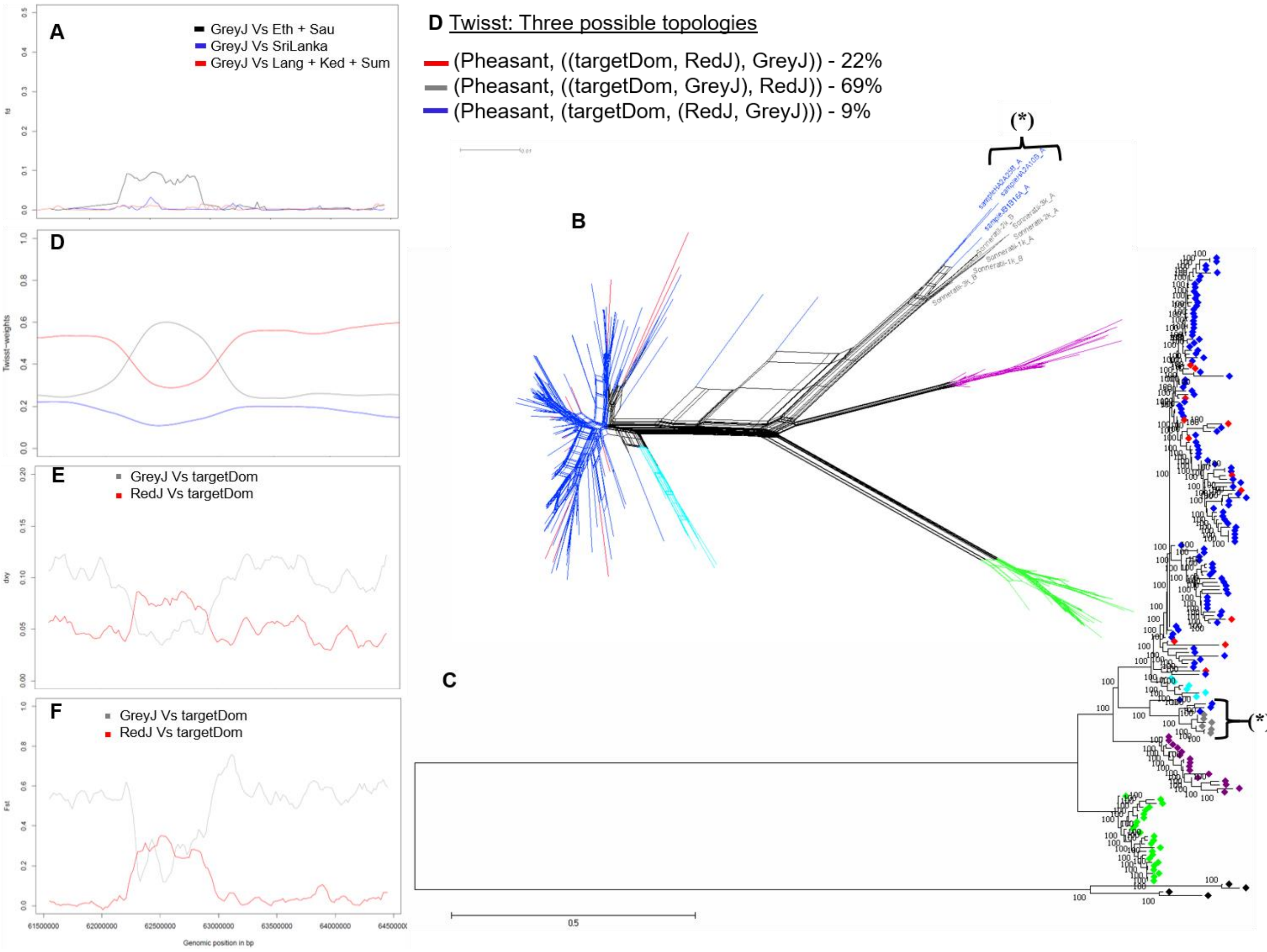


**Fig. S7.** A 200 kb (Chr 4: 62097304 - 62297319 bp) introgressed region from Grey junglefowl into domestic chicken.

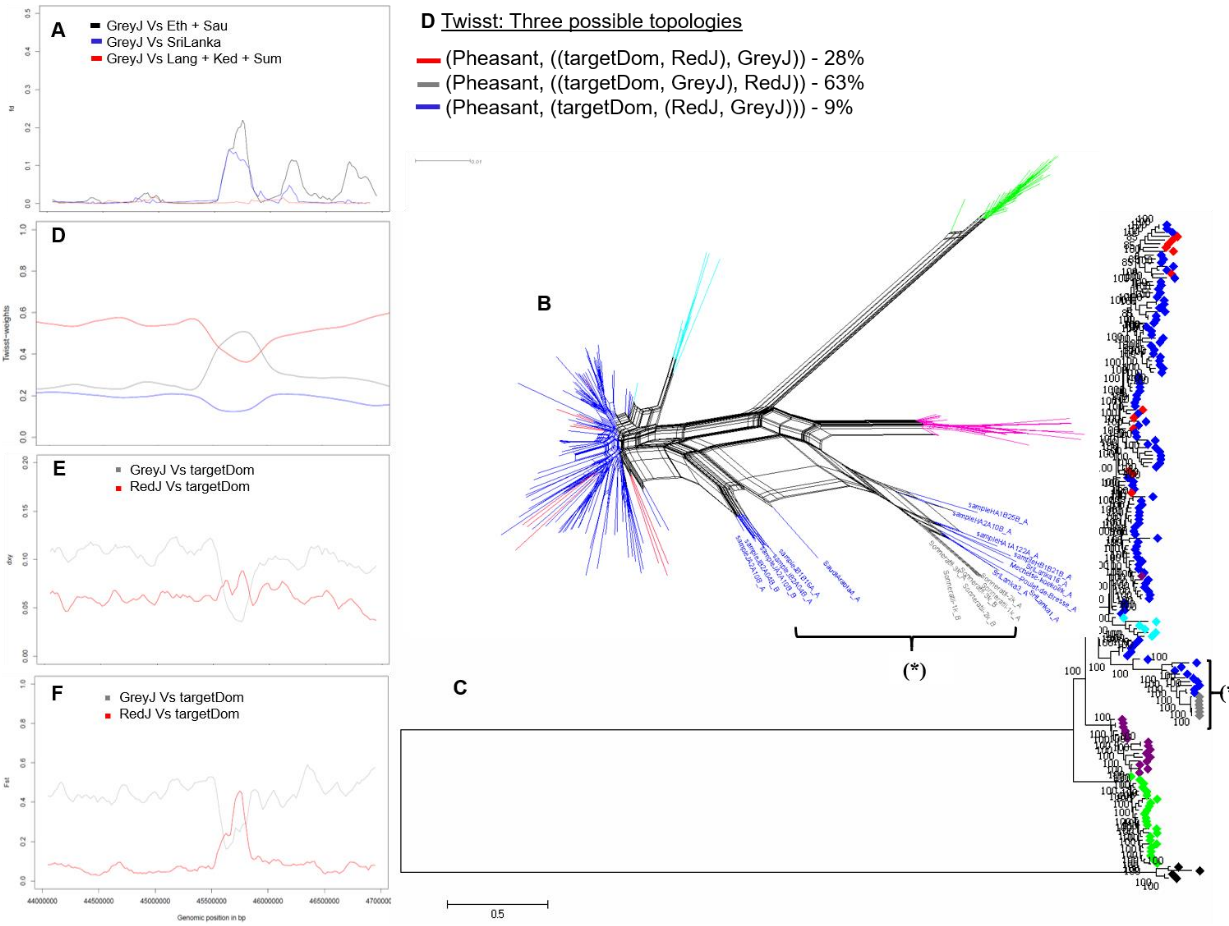


**Fig. S8.** A 280 kb (Chr 5: 45674368 - 45954418 bp) introgressed region from Grey junglefowl into domestic chicken.

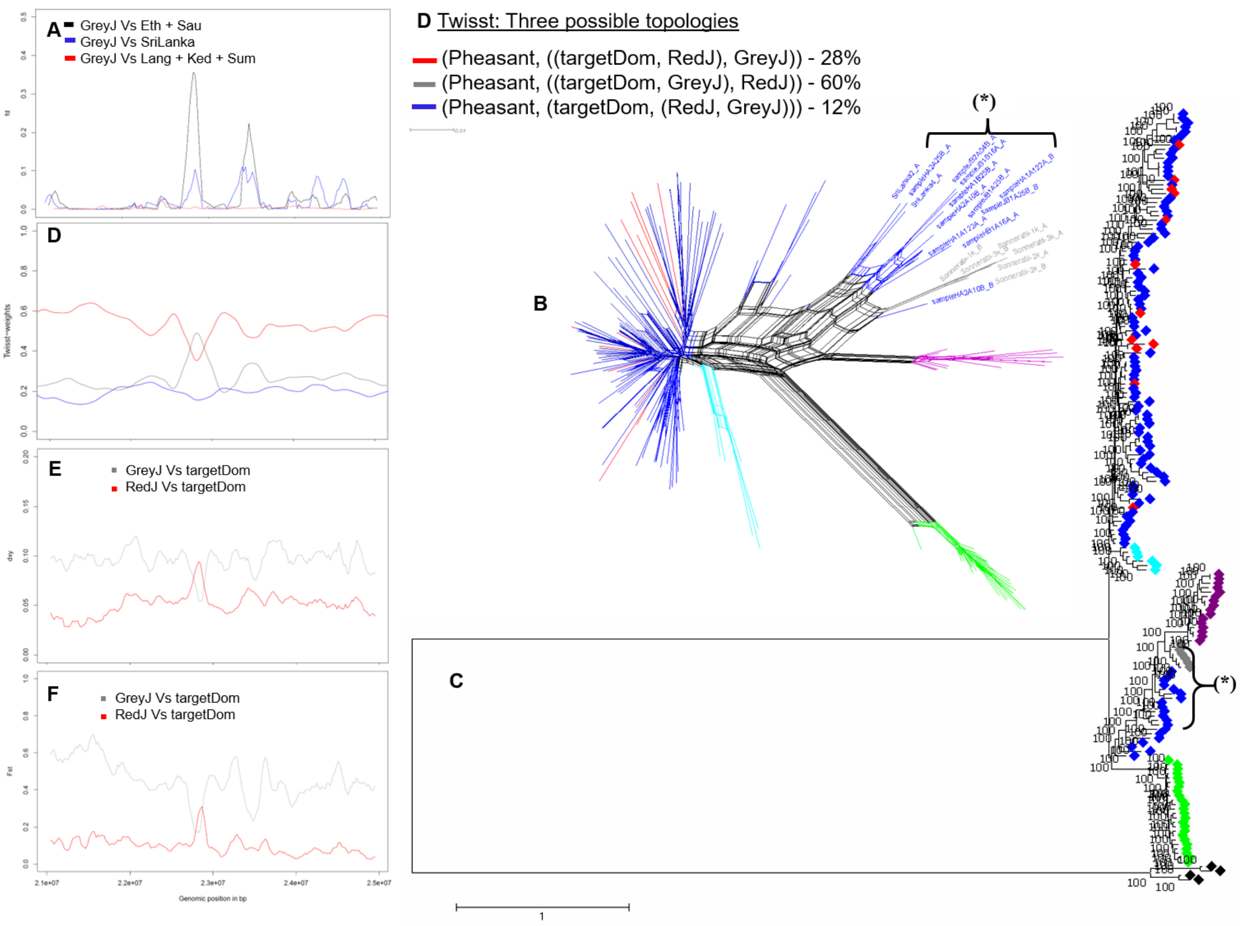


**Fig. S9.** A 140 kb (Chr 7: 22652767 - 22792759 bp) introgressed region from Grey junglefowl into domestic chicken.

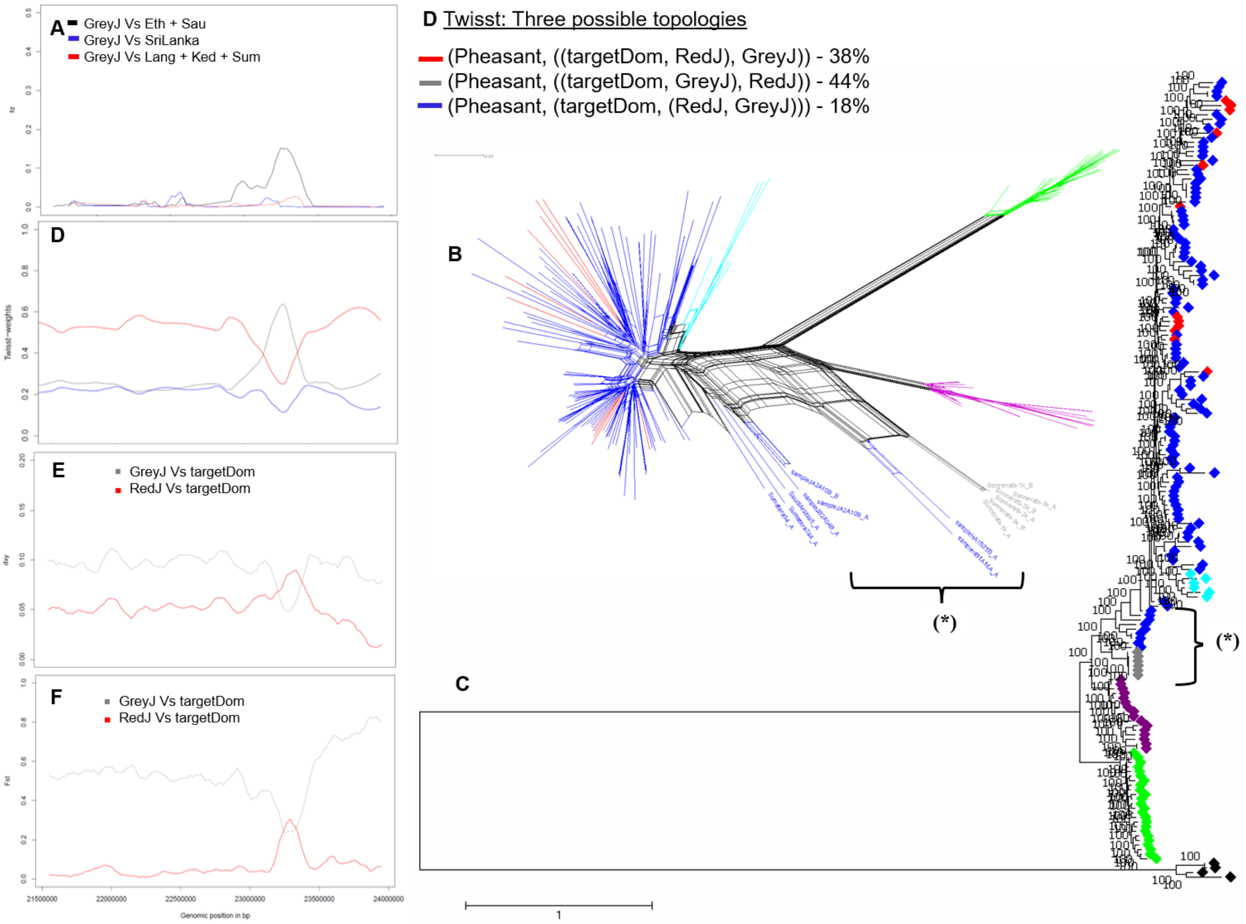


**Fig. S10.** A 500 kb (Chr 9: 23052049 - 23552045 bp) introgressed region from Grey junglefowl into domestic chicken.

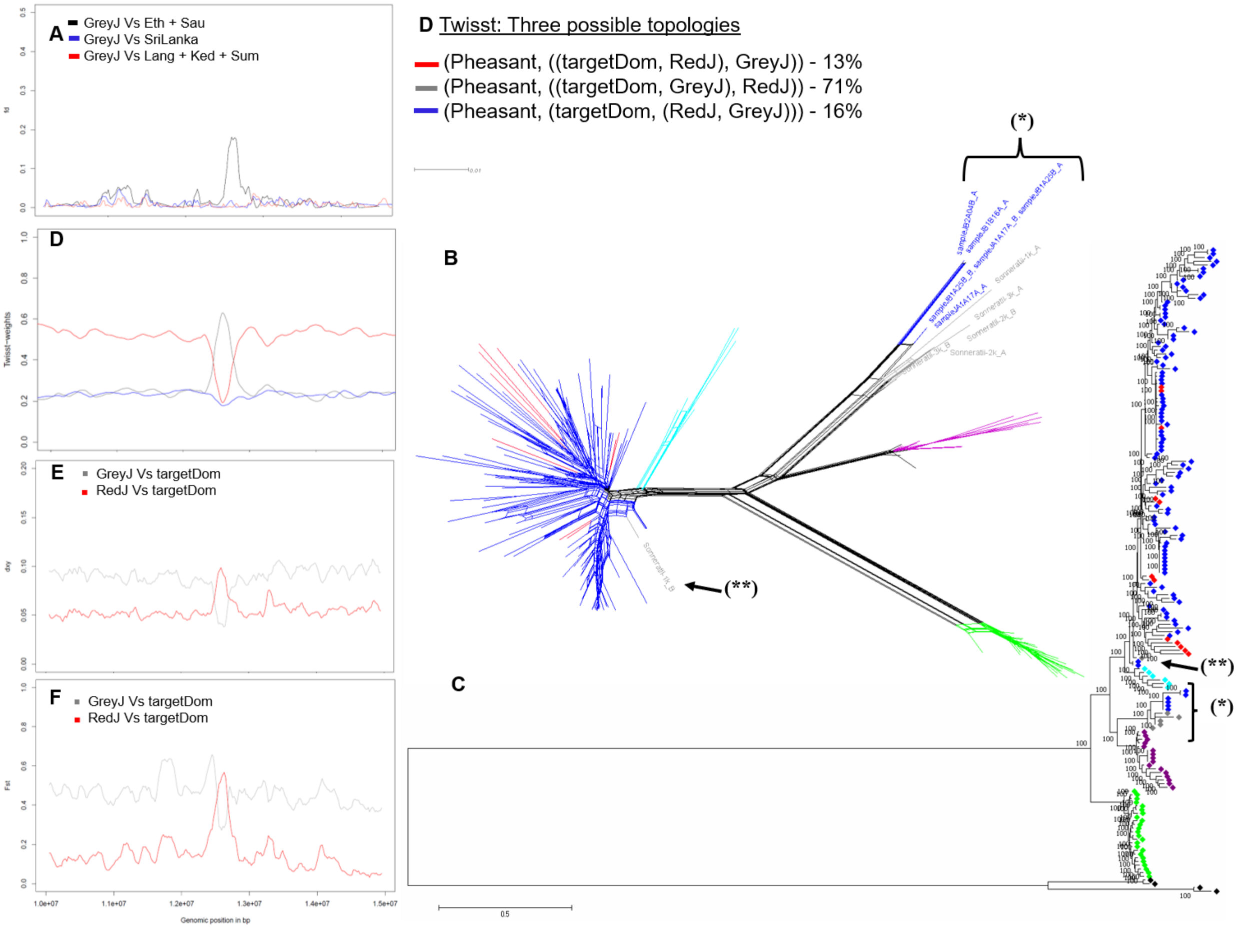


**Fig. S11.** A 100 kb (Chr 12: 12914268 - 13014266 bp) introgressed region from (*****) Grey junglefowl into domestic and (******) *vice versa*. The *Twisst* values and plots are based on the introgressed domestic haplotypes from the Grey junglefowl and does not account for the reverse introgression.

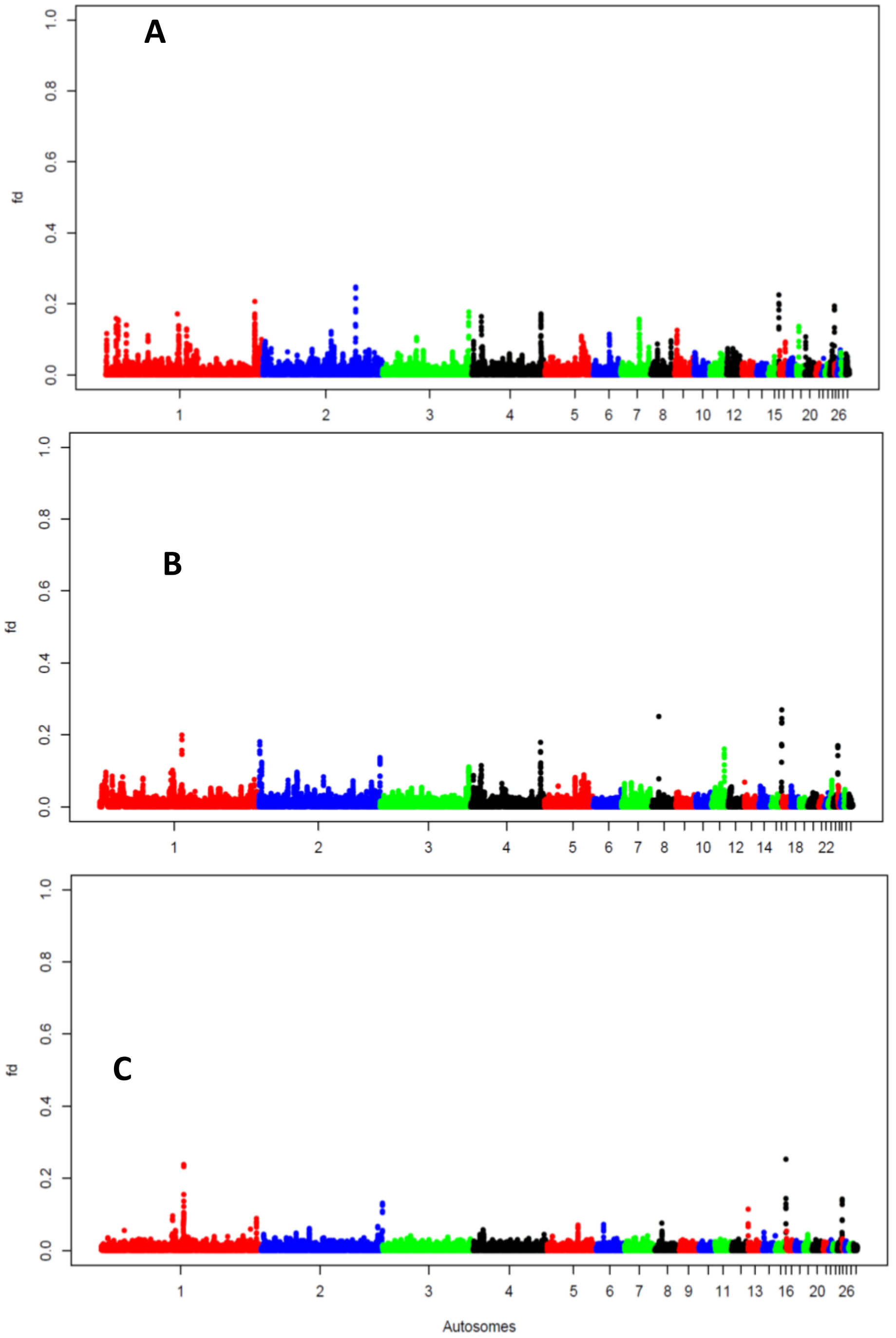


**Fig. S12.** The *fd* plots test for the comparison between Ceylon junglefowl and domestic chicken population from (**A**) Ethiopia and Saudi, (**B**) Sri Lanka and (**C**) Southeast and East Asia. The *Y*-axis fd value and X-axis 1 – 28 autosomes.

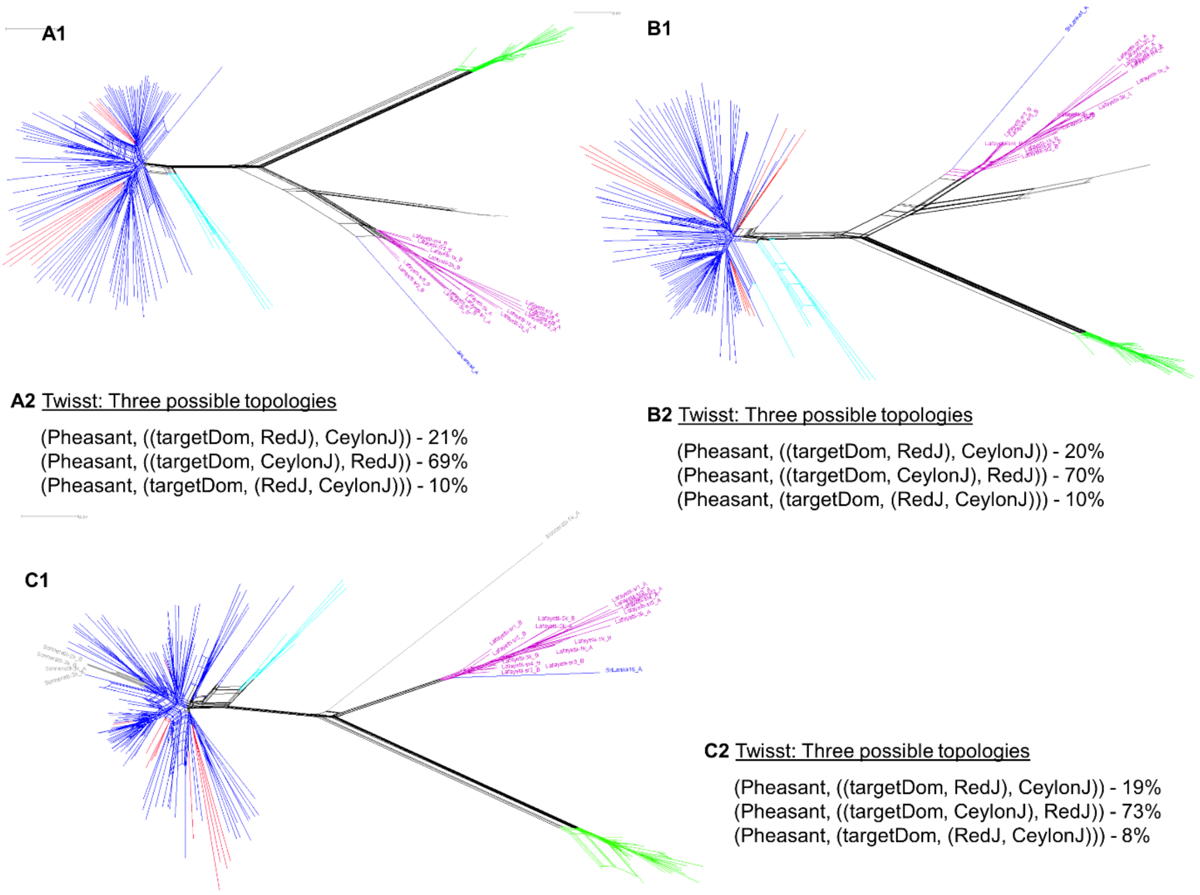


**Fig. S13.** Network and *Twisst* proportion of topologies for three Ceylon candidate introgressed regions into domestic chicken (**A - C**). (**A1**) and (**A2**) 6.52 Mb region Chr 1: 2895616 - 9418660 bp, (**B1**) and (**B2**) 3.95 Mb Chr 1: 25261354 - 29205161 bp, (**C1**) and (**C2**) 1.38 Mb region Chr 1: 147936229 - 149316591 bp. (**C1**) also shows support for introgression from domestic chicken to some Grey junglefowl haplotypes at the same region.

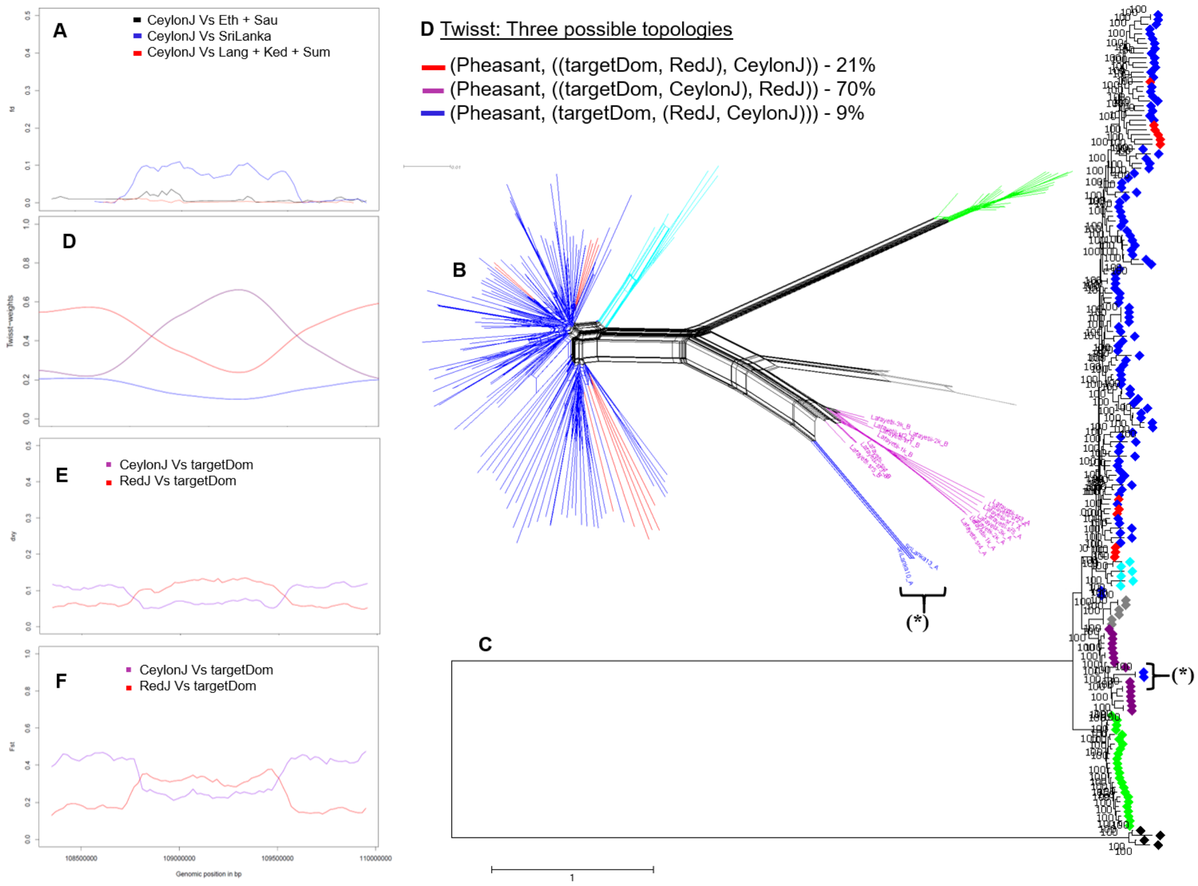


**Fig. S14.** A 600 kb (Chr 3: 108325801 - 108925723 bp) introgressed region from Ceylon junglefowl to domestic chicken.

(**A**) *fd* plot (**B**) haplotype-based network, (**C**) maximum likelihood tree, (**D**) *Twisst* plot and the proportion of each topology, (**E**) $D_{XY}$ and (**E**) $F_{ST}$. Eth, Sau, SriLanka, Lang, Ked, Sum are domestic chicken from Ethiopia, Saudi, Sri Lanka and Langshan, Kedu Hitam and Sumatra breeds, respectively, CeylonJ is Ceylon junglefowl and targetDom are the introgressed domestic chicken haplotypes (*****). ◆: Domestic chicken; ◆: Red junglefowl, ◆: Javan red junglefowl, ◆: Grey junglefowl, ◆: Ceylon junglefowl, ◆: Green junglefowl, ◆: common Pheasant.

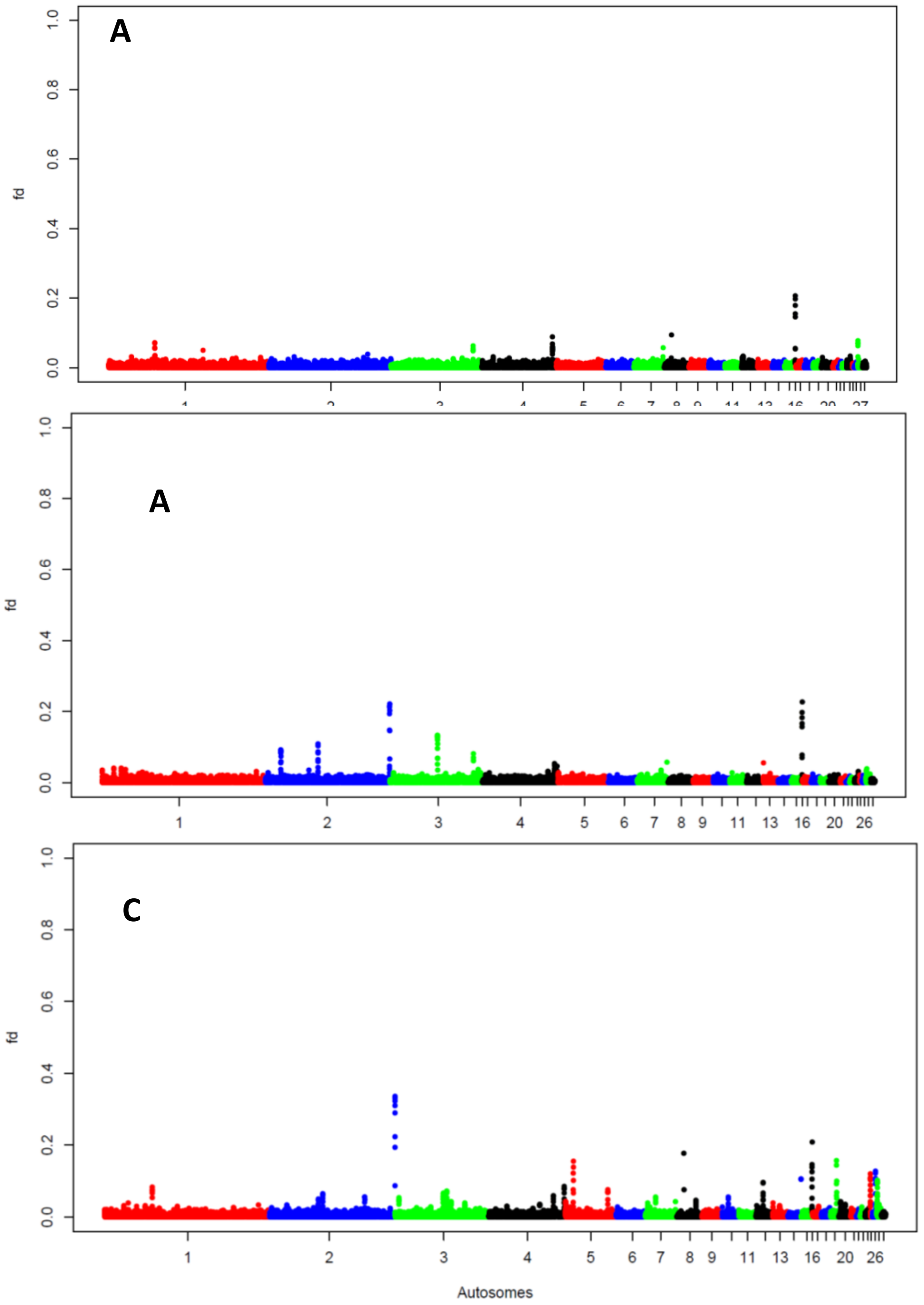


**Fig. S15.** The *fd* plots test for the comparison between Green junglefowl and domestic chicken population from (**A**) Ethiopia and Saudi, (**B**) Sri Lanka and (**C**) Southeast and East Asia. The Y-axis fd value and X-axis 1 – 28 autosomes.

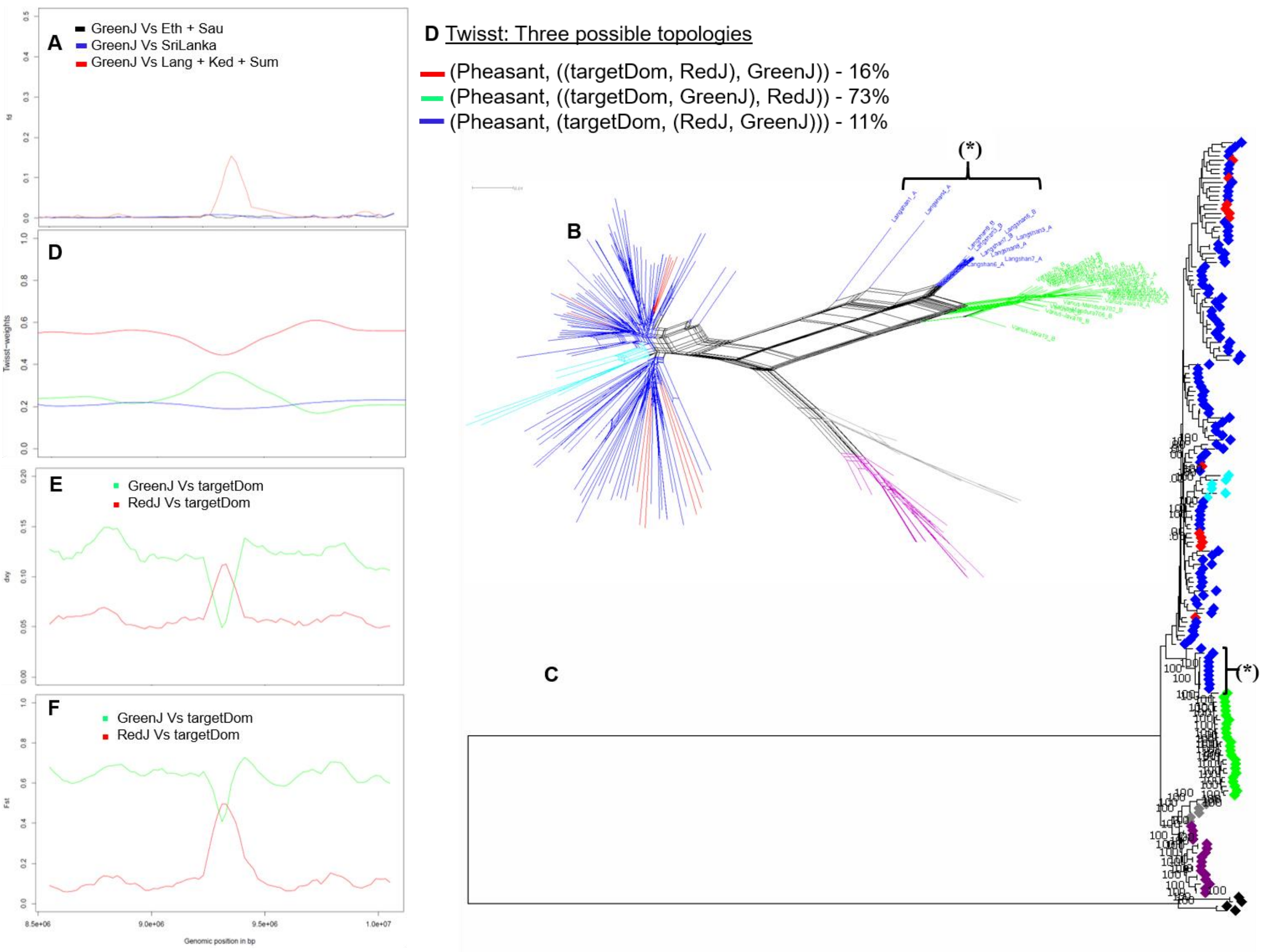


**Fig. S16:** A 100 kb (Chr 5: 9538715 - 9638713 bp) introgressed region from Green junglefowl into domestic chicken.

(**A**) *fd* plot, (**B**) haplotype-based network, (**C**) maximum likelihood tree, (**D**) TWISST plot and the proportion of each topology, (**E**) $D_{XY}$ and (**E**) $F_{ST}$. Eth, Sau, SriLanka, Lang, Ked, Sum are domestic chicken from Ethiopia, Saudi, Sri Lanka and Langshan, Kedu Hitam and Sumatra breeds, respectively, GreenJ is Green junglefowl and targetDom are the introgressed domestic haplotypes (*****). ◆: Domestic; ◆: Red junglefowl, ◆: Javan red junglefowl, ◆: Grey junglefowl, ◆: Ceylon junglefowl, ◆: Green junglefowl, ◆: common Pheasant.

**Table S1.** Sampling, mapping and variants statistics. HomAA and HetRA are the proportion of homozygous and heterozygous SNPs to the reference (*Galgal*5.0), respectively.

| Sample Name | Species Name | Geographic Distribution/Collection Point | Mapped Rate | Properly paired rate | HomAA% | HetRA% | Total SNPs (HomAA + HetRA) |
|---|---|---|---|---|---|---|---|
| HA1A12A | Gallus gallus domesticus | Ethiopia | 99.75 | 96.88 | 49.78 | 50.22 | 5496987 |
| HA1B25B | Gallus gallus domesticus | Ethiopia | 99.76 | 96.43 | 43.95 | 56.05 | 5715229 |
| HA2A10B | Gallus gallus domesticus | Ethiopia | 99.74 | 95.07 | 44.74 | 55.26 | 5702301 |
| HA2A25B | Gallus gallus domesticus | Ethiopia | 99.71 | 95.28 | 42.42 | 57.58 | 5772262 |
| HB1A16A | Gallus gallus domesticus | Ethiopia | 99.74 | 95.98 | 42.13 | 57.87 | 5818311 |
| HB1B21B | Gallus gallus domesticus | Ethiopia | 99.74 | 96.28 | 42.75 | 57.25 | 5779878 |
| JA1A17A | Gallus gallus domesticus | Ethiopia | 99.75 | 96.23 | 46.25 | 53.75 | 5611611 |
| JA2A10B | Gallus gallus domesticus | Ethiopia | 99.75 | 95.99 | 47.24 | 52.76 | 5577330 |
| JB1A25B | Gallus gallus domesticus | Ethiopia | 99.76 | 96.21 | 52.97 | 47.03 | 5345839 |
| JB1B16A | Gallus gallus domesticus | Ethiopia | 99.75 | 96.18 | 42.44 | 57.56 | 5764815 |
| JB2A04B | Gallus gallus domesticus | Ethiopia | 99.4 | 95.97 | 55.19 | 44.81 | 5303913 |
| SaudiArabia1 | Gallus gallus domesticus | Saudi Arabia | 99.53 | 93.43 | 51.31 | 48.69 | 5254695 |
| SaudiArabia2 | Gallus gallus domesticus | Saudi Arabia | 99.45 | 93.74 | 41.01 | 58.99 | 5685951 |
| SaudiArabia3 | Gallus gallus domesticus | Saudi Arabia | 99.07 | 94.8 | 44.38 | 55.62 | 5391149 |
| SaudiArabia4 | Gallus gallus domesticus | Saudi Arabia | 99.45 | 95.83 | 24.23 | 75.77 | 6540870 |
| SaudiArabia5 | Gallus gallus domesticus | Saudi Arabia | 99.16 | 93.96 | 41.35 | 58.65 | 5562922 |
| SriLanka1 | Gallus gallus domesticus | Sri Lanka | 99.14 | 94.21 | 36.75 | 63.25 | 6083602 |
| SriLanka10 | Gallus gallus domesticus | Sri Lanka | 99.1 | 94.6 | 37.4 | 62.6 | 6061984 |
| SriLanka12 | Gallus gallus domesticus | Sri Lanka | 99.21 | 93.83 | 35.27 | 64.73 | 6175086 |
| SriLanka13 | Gallus gallus domesticus | Sri Lanka | 99.15 | 93.42 | 37.41 | 62.59 | 6056414 |
| SriLanka15 | Gallus gallus domesticus | Sri Lanka | 99.16 | 93.1 | 36.98 | 63.02 | 6105444 |
| SriLanka16 | Gallus gallus domesticus | Sri Lanka | 99.12 | 94.37 | 36.27 | 63.73 | 6159000 |
| SriLanka2 | Gallus gallus domesticus | Sri Lanka | 99.04 | 94.38 | 35.84 | 64.16 | 6106133 |
| SriLanka3 | Gallus gallus domesticus | Sri Lanka | 99.21 | 94.58 | 36.76 | 63.24 | 6084710 |
| SriLanka4 | Gallus gallus domesticus | Sri Lanka | 98.98 | 94.51 | 35.08 | 64.92 | 6196012 |
| SriLanka5 | Gallus gallus domesticus | Sri Lanka | 99.23 | 94.38 | 35.81 | 64.19 | 6155422 |
| SriLanka9 | Gallus gallus domesticus | Sri Lanka | 99.2 | 94.27 | 34.83 | 65.17 | 6163149 |

| | | | | | | | |
|---|---|---|---|---|---|---|---|
| Langshan1 | Gallus gallus domesticus | Langshan, China - Collected in UK | 99.79 | 98.99 | 64.19 | 35.81 | 4183574 |
| Langshan2 | Gallus gallus domesticus | Langshan, China - Collected in UK | 99.89 | 99.07 | 62.76 | 37.24 | 4219311 |
| Langshan3 | Gallus gallus domesticus | Langshan, China - Collected in UK | 99.84 | 98.73 | 67.7 | 32.3 | 4052704 |
| Langshan4 | Gallus gallus domesticus | Langshan, China - Collected in UK | 99.87 | 98.3 | 59.37 | 40.63 | 4770695 |
| Langshan5 | Gallus gallus domesticus | Langshan, China - Collected in UK | 99.89 | 99.06 | 61.98 | 38.02 | 4090606 |
| Langshan6 | Gallus gallus domesticus | Langshan, China - Collected in UK | 99.86 | 99.03 | 61.29 | 38.71 | 4226957 |
| Langshan7 | Gallus gallus domesticus | Langshan, China - Collected in UK | 99.85 | 97.49 | 63.9 | 36.1 | 4499626 |
| Langshan8 | Gallus gallus domesticus | Langshan, China - Collected in UK | 99.87 | 98.29 | 62.04 | 37.96 | 4280925 |
| Sumatera738 | Gallus gallus domesticus | Indonesia | 99.14 | 98.04 | 50.58 | 49.42 | 4556806 |
| Sumatera741 | Gallus gallus domesticus | Indonesia | 99.12 | 97.52 | 57.44 | 42.56 | 4508322 |
| Sumatera744 | Gallus gallus domesticus | Indonesia | 99.1 | 97.99 | 58.61 | 41.39 | 4450663 |
| Sumatera757 | Gallus gallus domesticus | Indonesia | 99.1 | 97.26 | 51.82 | 48.18 | 4659515 |
| Sumatera94 | Gallus gallus domesticus | Indonesia | 99.05 | 97.13 | 55.83 | 44.17 | 4193805 |
| KeduHitam761 | Gallus gallus domesticus | Indonesia | 99.62 | 98.56 | 43.22 | 56.78 | 5458556 |
| KeduHitam763 | Gallus gallus domesticus | Indonesia | 99.59 | 98.48 | 42.22 | 57.78 | 5551599 |
| KeduHitam766 | Gallus gallus domesticus | Indonesia | 99.63 | 98.61 | 53.71 | 46.29 | 5205120 |
| KeduHitam767 | Gallus gallus domesticus | Indonesia | 99.56 | 98.29 | 40.28 | 59.72 | 5745786 |
| KeduHitam778 | Gallus gallus domesticus | Indonesia | 98.94 | 97.66 | 54.11 | 45.89 | 4996704 |
| KeduHitam779 | Gallus gallus domesticus | Indonesia | 99.3 | 97.61 | 44.14 | 55.86 | 5324021 |
| KeduHitam780 | Gallus gallus domesticus | Indonesia | 99.5 | 97.92 | 45.98 | 54.02 | 5424436 |
| KeduHitam781 | Gallus gallus domesticus | Indonesia | 99.06 | 98 | 56.45 | 43.55 | 5035441 |
| KeduHitam783 | Gallus gallus domesticus | Indonesia | 99.05 | 97.81 | 46.95 | 53.05 | 5126861 |
| KeduHitam784 | Gallus gallus domesticus | Indonesia | 99.52 | 98.35 | 44.98 | 55.02 | 5238868 |
| Mechelse-koekoek | Gallus gallus domesticus | Europe | 96.9 | 91.9 | 55.47 | 44.53 | 5206378 |
| Mechelse-styrian | Gallus gallus domesticus | Europe | 97 | 92.89 | 34.66 | 65.34 | 6163592 |
| Poulet-de-Bresse | Gallus gallus domesticus | Europe | 97.63 | 93.32 | 47.83 | 52.17 | 5554109 |
| RedJunglefowl1 | Gallus gallus (Red junglefowl) | Southeast Asia | 98.69 | 93.85 | 38.55 | 61.45 | 6090522 |
| RedJunglefowl2 | Gallus gallus (Red junglefowl) | Southeast Asia | 99.53 | 96.65 | 38.48 | 61.52 | 6322049 |
| RedJunglefowl3 | Gallus gallus (Red junglefowl) | Southeast Asia | 99.52 | 95.25 | 37.1 | 62.9 | 6721117 |
| RedJunglefowl4 | Gallus gallus (Red junglefowl) | Southeast Asia | 99.51 | 96.66 | 43.85 | 56.15 | 6432497 |
| RedJunglefowl5 | Gallus gallus (Red junglefowl) | Southeast Asia | 99.32 | 96 | 34.17 | 65.83 | 7055675 |

| | | | | | | | |
|---|---|---|---|---|---|---|---|
| RedJungleFowlKoen | Gallus gallus (Red junglefowl) | South/Southeast Asia - Private collection | 95.66 | 92.14 | 33.62 | 66.38 | 6790156 |
| Bankiva650 | Gallus gallus bankiva (Javan red junglefowl) | Indonesia | 98.85 | 97.46 | 60.09 | 39.91 | 6880685 |
| Bankiva759 | Gallus gallus bankiva (Javan red junglefowl) | Indonesia | 98.98 | 97.57 | 59.49 | 40.51 | 6890215 |
| Bankiva760 | Gallus gallus bankiva (Javan red junglefowl) | Indonesia | 98.87 | 97.16 | 60.1 | 39.9 | 6811363 |
| Sonneratii1k | Gallus sonneratii (Grey junglefowl) | Indian subcontinent - Private collection | 99.69 | 95.4 | 77.76 | 22.24 | 10050824 |
| Sonneratii2k | Gallus sonneratii (Grey junglefowl) | Indian subcontinent - Private collection | 99.61 | 88.02 | 75.64 | 24.36 | 10115589 |
| Sonneratii3k | Gallus sonneratii (Grey junglefowl) | Indian subcontinent - Private collection | 99.69 | 95.4 | 84.74 | 15.26 | 9432160 |
| Lafayetii1k | Gallus lafayetti (Ceylon junglefowl) | Sri Lanka - Private collection | 99.61 | 90.84 | 79.78 | 20.22 | 10637322 |
| Lafayetii2k | Gallus lafayetti (Ceylon junglefowl) | Sri Lanka - Private collection | 99.66 | 93.6 | 79.84 | 20.16 | 10619033 |
| Lafayetii3k | Gallus lafayetti (Ceylon junglefowl) | Sri Lanka - Private collection | 99.67 | 95.1 | 85.73 | 14.27 | 10267990 |
| Lafayetiisr1 | Gallus lafayetti (Ceylon junglefowl) | Sri Lanka - Private collection | 98.93 | 92.66 | 74.06 | 25.94 | 11126055 |
| Lafayetiisr2 | Gallus lafayetti (Ceylon junglefowl) | Sri Lanka | 99.05 | 91.09 | 74.69 | 25.31 | 11091506 |
| Lafayetiisr3 | Gallus lafayetti (Ceylon junglefowl) | Sri Lanka | 98.88 | 91.5 | 72.73 | 27.27 | 11216350 |
| Lafayetiisr4 | Gallus lafayetti (Ceylon junglefowl) | Sri Lanka | 98.98 | 92.83 | 72.7 | 27.3 | 11215158 |
| Lafayetiisr5 | Gallus lafayetti (Ceylon junglefowl) | Sri Lanka | 98.75 | 89.28 | 72.53 | 27.47 | 11218503 |
| Varius1k | Gallus varius (Green junglefowl) | Indonesia - Private collection | 99.52 | 85.65 | 87.94 | 12.06 | 12019627 |
| Varius2k | Gallus varius (Green junglefowl) | Indonesia - Private collection | 99.54 | 90.03 | 88.04 | 11.96 | 12004164 |
| Varius3k | Gallus varius (Green junglefowl) | Indonesia - Private collection | 99.61 | 94.6 | 86.74 | 13.26 | 11958988 |
| VariusJava18 | Gallus varius (Green junglefowl) | Indonesia | 98.54 | 96.26 | 90.56 | 9.44 | 10591696 |
| VariusJava19 | Gallus varius (Green junglefowl) | Indonesia | 98.47 | 96.51 | 82.54 | 17.46 | 11112951 |
| VariusMandura702 | Gallus varius (Green junglefowl) | Indonesia | 98.56 | 95.01 | 92.29 | 7.71 | 11077799 |
| VariusMandura703 | Gallus varius (Green junglefowl) | Indonesia | 98.83 | 94.31 | 88 | 12 | 11590408 |
| VariusMandura704 | Gallus varius (Green junglefowl) | Indonesia | 98.77 | 94.76 | 91.51 | 8.49 | 10841620 |
| VariusMandura706 | Gallus varius (Green junglefowl) | Indonesia | 98.71 | 93.73 | 89.48 | 10.52 | 11414585 |
| VariusMandura710 | Gallus varius (Green junglefowl) | Indonesia | 98.76 | 95.26 | 90.96 | 9.04 | 10429506 |

| | | | | | | | |
|---|---|---|---|---|---|---|---|
| VariusMandura714 | Gallus varius (Green junglefowl) | Indonesia | 98.8 | 94.82 | 92.26 | 7.74 | 10620997 |
| VariusMandura715 | Gallus varius (Green junglefowl) | Indonesia | 98.69 | 95.02 | 90.07 | 9.93 | 11322992 |
| PheasantsFemale | Phasianus colchicus | Collected in UK | 91.95 | 81.69 | 96.01 | 3.99 | 53278629 |
| PheasantsMale | Phasianus colchicus | Collected in UK | 93 | 83.47 | 94.19 | 5.81 | 56157598 |

**Table S2*A***. Candidate introgressed regions from domestic chicken and/or Red junglefowl into Grey/Ceylon junglefowls

| Candidate introgressed regions* | Length | Genes within the candidate introgressed regions*** |
|---|---|---|
| ***Domestic chicken/Red junglefowl into Grey Junglefowl*** | | |
| Chr1: 141.0 – 167.0 | 26 Mb | *TNFSF13B, ABHD13, LIG4, FAM155A, ARGLU1, EFNB2, SLC10A2, ERCC5, BIVM, KDELC1, TEX30, METTL21C, TPP2, FHF-4, ITGBL1, NALCN, TMTC4, GGACT, PCCA, ZIC2, ZIC5, CLYBL, TM9SF2, gga-mir-2984, UBAC2, GPR183, GPR18, DOCK9, SLC15A1, STK24, FARP1, gga-mir-1555, IPO5, RAP2A, MBNL2, UGGT2, DNAJC3, DZIP1, CLDN10, ABCC4, SOX21, GPR180, TGDS, DCT, RF00066, GPC5, gga-mir-92-1, gga-mir-19b, gga-mir-20a, gga-mir-19a, gga-mir-18a, gga-mir-17, SLITRK6, SLITRK1, SPRY2, NDFIP2, RBM26, RNF219, POU4F1, EDNRB, SLAIN1, MYCBP2, FBXL3, CLN5, GATD3A, ACOD1, KCTD12, LMO7, UCHL3, TBC1D4, KLF12, KLF5, PIBF1, DIS3, BORA, MZT1, DACH1, gga-mir-1743, KLHL1, PCDH9, RF00154, gga-mir-7445-2, TDRD3, DIAPH3, PCDH17, RF02271, RF00493, RF00494, OLFM4, PCDH8, CNMD, SUGT1, ELF1, WBP4, MTRF1, RGCC, VWA8* |
| Chr2: 11.0 – 20.0 | 9 Mb | *PFKP, PITRM1, KLF6, gga-mir-6628, GJD4, CCNY, CREM, CUL2, PARD3, RF02271, NRP1, ITGB1, EPC1, gga-mir-1768, KIF5B, ARHGAP12, ZEB1, ZNF438, SVIL, JCAD, MTPAP, MAP3K8, BAMBI, WAC, MPP7, ARMC4, MKX, RAB18, YME1L1, MASTL, ACBD5, ABI1, PDSS1, APBB1IP, GAD2, MYO3A, GPR158, THNSL1, ENKUR, PRTFDC1, ARHGAP21, KIAA1217, PTF1A, ARMC3, PIP4K2A, SPAG6, gga-mir-12240, BMI1, COMMD3, DNAJC1, MLLT10, RF00001, NEBL, PLXDC2, MALRD1, ARL5B, NSUN6, CACNB2, SLC39A12, MMR1L2, MMR1L1, MMR1L3, MMR1L4, MRC1, STAM, HACD1, VIM, ST8SIA6, gga-mir-1661, TRDMT1, CUBN* |
| Chr4:76.4 – 79.2 | 2.8 Mb | *TAPT1, Prom1, FGFBP2, CD38, BST1, FBXL5, CC2D2A, C1QTNF7, CPEB2, NKX3-2, RAB28, HS3ST1, ZNF518B, WDR1, SLC2A9, DRD5, OTOP1, TMEM128, LYAR, ZBTB49, NSG1, STX18, MSX1, CYTL1, STK32B, EVC2, EVC, CRMP1B* |
| ***Domestic chicken into Ceylon junglefowl*** | | |
| Chr5: 49.33 – 49.43 | 100 kb | *No gene* |

*Positions along the chromosome in megabase (Mb),

**Table S2*B***. Candidate introgressed regions from non-red junglefowls into domestic chicken/Red junglefowl

| **Candidate introgressed regions*** | **Length** | **Proportion of haplotypes introgressed in each population (%)**** | | | | | | | **Genes within the candidate introgressed regions***** |
|---|---|---|---|---|---|---|---|---|---|
| | | Ethiopia (n = 22) | Saudi Arabia (n = 10) | Sri Lanka (n = 22) | SEA Langshan (n = 16) | SEA Kedu Hitam (n = 20) | SEA Sumatra (n = 10) | Red Junglefowl (n = 12) | |
| ***Introgressed regions from Grey junglefowl to domestic chicken/Red junglefowl*** | | | | | | | | | |
| **Chr2:119.68** – 119.90 | 220 kb | 23 | 20 | 9 | 0 | 0 | 0 | 8 | - |
| Chr3:50.75 – 50.85 | 100 kb | 27 | 0 | 23 | 0 | 5 | 10 | 0 | *NOX3* |
| Chr4:62.10 – 62.30 | 200 kb | 14 | 0 | 0 | 0 | 0 | 0 | 0 | *RF00003* |
| Chr5:45.67 – 45.95 | 280 kb | 41 | 10 | 9 | 0 | 0 | 0 | 0 | *PPP4R4*, *SERPINA10*, *SPIA4*, *SPIA1*, *GSC* |
| Chr6:20.72 – 20.84 | 120 kb | 45 | 0 | 0 | 0 | 0 | 0 | 0 | *IDE*, *Mar-05*, *CPEB3* |
| Chr7:22.65 – 22.79 | 140 kb | 50 | 0 | 9 | 0 | 0 | 0 | 0 | - |
| Chr9:23.05 – 23.55 | 500 kb | 23 | 10 | | 0 | 0 | 20 | 0 | *KCNAB3*, *GMPS*, *gga-mir-1658*, *C3orf33*, *PLCH1*, *MME*, *GPR149*, *DHX36*, *RAP2B*, *ARHGEF26*, *P2RY1*, *MBNL1* |
| Chr12: 12.54 – 12.64 | 100 kb | 27 | 0 | 0 | 0 | 0 | 0 | 0 | *FHIT* |
| ***Introgressed regions from Ceylon junglefowl to domestic chicken*** | | | | | | | | | |
| **Chr1:2.90 – 9.42** | 6.52 Mb | 0 | 0 | 5 | 0 | 0 | 0 | 0 | *PLXNA4*, *gga-mir-6621*, *PODXL*, *MKLN1*, *gga-mir-29b-1*, *gga-mir-29a*, *K123*, *IL2RA*, *RBM17*, *PFKFB3*, *SFMBT2*, *ITIH5*, *ITIH2*, *KIN*, *ATP5F1C*, *TAF3*, *GATA3*, *CELF2*, *gga-mir-1626*, *gga-mir-1596*, *USP6NL*, *ECHDC3*, *UPF2*, *DHTKD1*, *SEC61A2*, *NUDT5*, *CDC123*, *CAMK1D*, *CCDC3*, *OPTN*, *MCM10*, *PHYH*, *SEPHS2L*, *BEND7*, *FRMD4A*, *gga-mir-1460*, *FAM107B*, *HSPA14*, *SUV39H2*, *DCLRE1C*, *MEIG1*, *TMEM243*, *DMTF1*, *RF02271*, *KIAA1324L*, *GRM3*, *SEMA3D*, *SEMA3A* |
| Chr1:25.25 – 29.21 | **3.95** Mb | 0 | 0 | 5 | 0 | 0 | 0 | 0 | *TES*, *TFEC*, *MDFIC*, *FOXP2*, *PPP1R3A*, *GPR85*, *gga-mir-1695*, *BMT2*, *TMEM168*, *LSMEM1*, *IFRD1*, *ZNF277*, *DOCK4*, *IMMP2L*, *LRRN3*, |

| | | | | | | | | | |
|---|---|---|---|---|---|---|---|---|---|
| | | | | | | | | | *DNAJB9, THAP5, AVPR2, PNPLA8, NRCAM, gga-mir-12208, CNTN1, PDZRN4* |
| Chr1:147.94 – 149.32 | 1.38 Mb | 0 | 0 | 5 | 0 | 0 | 0 | 0 | *GPC6, RF00066, GPC5* |
| Chr3:108.33 – 108.93 | 600 kb | 0 | 0 | 9 | 0 | 0 | 0 | 0 | *CRISP3, CRISP2, RHAG, CYP2AC1, CYP2AC2, CENPQ, MMUT, OPN5L2, FOXP2* |
| ***Introgressed region from Green junglefowl to domestic chicken*** | | | | | | | | | |
| Chr5: 9.54 – 9.64 | 100 kb | 0 | 0 | 0 | 63 | 0 | 0 | 0 | *SWAP70, WEE1, ZNF143, IPO7, RF00319* |

*Positions along the chromosome in megabase (Mb), **SEA (South-East and East Asia), ***Ensembl release version 96

**Table S3.** Functional annotations for the enriched genes within the introgressed regions

| **Gene ontology for introgressed genes from domestic chicken to Grey junglefowl** | | | |
|---|---|---|---|
| **Term** | **Biological function** | **P-Value** | **Genes** |
| GO:0002042 | cell migration involved in sprouting angiogenesis | 0.003885481 | *NRP1*, *EFNB2*, *ITGB1* |
| GO:0030890 | positive regulation of B cell proliferation | 0.004098509 | *BMI1*, *GPR183*, *CD38*, *BST1* |
| GO:0005886 | plasma membrane | 0.004886068 | *GPR183*, *RAP2A*, *SLC2A9*, *GPR158*, *OLFM4*, *NRP1*, *BST1*, *VIM*, *EFNB2*, *PCDH9*, *CLDN10*, *PCDH8*, *LIG4*, *PCDH17*, *APBB1IP*, *WBP4*, *ITGB1*, *CD38*, *GPC5*, *SVIL*, *NALCN*, *BAMBI*, *PIP4K2A*, *DNAJC1* |
| GO:0007224 | smoothened signaling pathway | 0.015997843 | *EVC2*, *EVC*, *DZIP1*, *CC2D2A* |
| GO:0046872 | metal ion binding | 0.01615502 | *DCT*, *ZBTB49*, *KLF6*, *NRP1*, *ZNF518B*, *KLF12*, *DZIP1*, *MBNL2*, *LIG4*, *ZEB1*, *ITGB1*, *ZNF438*, *PCCA*, *RBM26* |
| GO:0045595 | regulation of cell differentiation | 0.017578595 | *SPRY2*, *SOX21*, *STK24* |
| GO:0003953 | NAD+ nucleosidase activity | 0.022841738 | *CD38*, *BST1* |
| GO:0001227 | transcriptional repressor activity, RNA polymerase II transcription regulatory region sequence-specific binding | 0.033003736 | *MSX1*, *KLF12*, *ZEB1* |
| GO:0045814 | negative regulation of gene expression, epigenetic | 0.047610409 | *BMI1*, *EPC1* |
| **Gene ontology for introgressed genes from Grey junglefowl to domestic chicken** | | | |
| GO:0050435 | Beta-amyloid metabolic process | 0.009746297 | *IDE*, *MME* |
| GO:0003725 | Double-stranded RNA binding | 0.044235391 | *DHX36*, *MBNL1* |
| **Gene ontology for introgressed genes from Ceylon junglefowl to domestic chicken** | | | |
| GO:0030212 | hyaluronan metabolic process | 0.021181 | *ITIH5*, *ITIH2* |
| GO:0009791 | Post-embryonic development | 0.025842519 | *MUT*, *GATA3*, *FOXP2* |
| GO:0000380 | alternative mRNA splicing, via spliceosome | 0.036777 | *CELF2*, *RBM17* |
| GO:0045786 | negative regulation of cell cycle | 0.04704 | *GATA3*, *THAP5* |
| GO:0005634 | nucleus | 0.039752 | *TAF3*, *CENPQ*, *THAP5*, *CDC5L*, *OPTN*, *KIN*, *SUV39H2*, *FOXP2*, *DMTF1*, *MDFIC*, *TFEC*, *GATA3*, *CELF2*, *IFRD1*, *CAMK1D*, *TES* |
| GO:0001135 | transcription factor activity, RNA polymerase II transcription factor recruiting | 0.019549 | *DMTF1*, *CDC5L* |
| GO:0038191 | neuropilin binding | 0.048175 | *SEMA3D*, *SEMA3A* |
| GO:0044212 | transcription regulatory region DNA binding | 0.049902 | *DMTF1*, *GATA3*, *CDC5L* |

3
